# A Population Receptive Field Model of the Magnetoencephalography Response

**DOI:** 10.1101/2020.08.28.272534

**Authors:** Eline R Kupers, Akhil Edadan, Noah C Benson, Wietske Zuiderbaan, Maartje C de Jong, Serge O Dumoulin, Jonathan Winawer

**Author notes:** indicates shared first authorship (Eline R Kupers, Akhil Edadan). indicates shared senior authorship (Serge O Dumoulin, Jonathan Winawer).

## Abstract

Computational models which predict the neurophysiological response from experimental stimuli have played an important role in human neuroimaging. One type of computational model, the population receptive field (pRF), has been used to describe cortical responses at the millimeter scale using functional magnetic resonance imaging (fMRI) and electrocorticography (ECoG). However, pRF models are not widely used for non-invasive electromagnetic field measurements (EEG/MEG), because individual sensors pool responses originating from several centimeter of cortex, containing neural populations with widely varying spatial tuning. Here, we introduce a forward-modeling approach in which pRFs estimated from fMRI data are used to predict MEG sensor responses. Subjects viewed contrast-reversing bar stimuli sweeping across the visual field in separate fMRI and MEG sessions. Individual subject’s pRFs were modeled on the cortical surface at the millimeter scale using the fMRI data. We then predicted cortical time series and projected these predictions to MEG sensors using a biophysical MEG forward model, accounting for the pooling across cortex. We compared the predicted MEG responses to observed visually evoked steady-state responses measured in the MEG session. We found that pRF parameters estimated by fMRI could explain a substantial fraction of the variance in steady-state MEG sensor responses (up to 60% in individual sensors). Control analyses in which we artificially perturbed either pRF size or pRF position reduced MEG prediction accuracy, indicating that MEG data are sensitive to pRF properties derived from fMRI. Our model provides a quantitative approach to link fMRI and MEG measurements, thereby enabling advances in our understanding of spatiotemporal dynamics in human visual field maps.

## 2 Introduction

A fundamental goal in human neuroscience is to understand how sensory inputs are transformed and represented in the nervous system. One approach to reach this goal is to build encoding models. This approach uses a quantitative description of the operations that relate input to output, *e.g.,* a visual image to fMRI blood-oxygen-level-dependent (BOLD) responses, providing a test of our understanding of how visual inputs are encoded in the visual pathways (Naselaris, Kay, Nishimoto, & Gallant, 2011; Holdgraf et al., 2017). Encoding models have been successful in predicting neural responses in human visual cortex. For example, visual field preferences of neural populations were predicted from fMRI BOLD responses (Dumoulin & Wandell, 2008; Kay, Winawer, Mezer, & Wandell, 2013) and intracranial field potentials, or electrocorticography (ECoG) (Yoshor, Bosking, Ghose, & Maunsell, 2007; Harvey, Vansteensel, et al., 2013; Winawer et al., 2013). In addition to providing a functional description of neural processes, encoding models can be used to compare data across different measurement techniques. For example, the fMRI BOLD signal measures vascular responses on the time scale of hundreds of milliseconds to seconds, whereas MEG measures magnetic flux at the millisecond scale; the data are not directly comparable but by applying a common encoding model with stimulus-referred parameters, such as position or size of the receptive field, the measurements can be compared. In this way, there is greater potential to integrate recordings with a high spatial resolution and recordings with a high temporal resolution, in order to study the visual system with greater precision.

However, encoding models from stimulus to measurement are relatively uncommon for non-invasive electromagnetic field measurements, like magnetoencephalography (MEG) or electroencephalography (EEG). While both MEG and EEG are widely used and provide excellent time-resolved measurements of brain activity across the whole brain, the pooling area of a single EEG or MEG sensor spans large parts of the cortex (on the order of several centimeters). Since this pooling area is much coarser than the spatial scale at which stimulus-selectivity tends to vary in visual cortex (on the order of millimeters for stimulus position, and sub-millimeter for orientation, spatial frequency, and other properties), building an encoding model to fit data from an MEG or EEG sensor is not straightforward, and may not be easily interpretable. For example, a population receptive field (pRF) for a single MEG sensor is likely to reflect neural signals from many different parts of the visual field and from multiple visual areas. The other way around, estimating local pRFs on the cortex from MEG sensor responses would require a computational model that transforms magnetic flux from hundreds of sensors to thousands of cortical locations. This inverse problem is ill-posed (under-constrained) and hence does not have a unique solution.

Here, we propose a novel, pRF modeling approach to predict MEG sensor responses from the stimulus. To do so, we extend the pRF model developed by Dumoulin and Wandell (2008), which has been a well-established approach to study the spatial properties of the human visual system in both healthy and diseased subjects (Wandell & Winawer, 2015; Dumoulin & Knapen, 2018). Our modeling approach can be divided into two steps. First, it estimates local pRFs on the cortex using fMRI, and predicts the neural response for a particular visual stimulus on the cortical surface. Second, the model projects these predicted responses to MEG sensors, using a biophysical model of the head. We compared predicted MEG sensor responses to observed MEG responses while subjects viewed a visual mapping stimulus. Using this modeling approach, we show that MEG responses to a visual stimulus can be predicted using pRF models estimated from fMRI.

## 3 Methods

### 3.1 Subjects

Ten subjects (5 female), ages 20-45 years (M = 29.7 years, SD = 7.3 years) with normal or correct-to-normal vision, participated in the study. MRI and MEG sessions were conducted on separate days. All scanning sessions took place at New York University. Subjects provided written informed consent. The experimental protocol was in compliance with the safety guidelines for MRI and MEG research and was approved by the University Committee on Activities involving Human Subjects at New York University, USA.

### 3.2 Stimuli

Stimuli were generated using MATLAB (MathWorks, MA, USA) and PsychToolbox (Brainard, 1997; Pelli, 1997; Kleiner et al., 2007) on a Macintosh computer. In both MRI and MEG sessions, subjects were presented with contrast-reversing checkerboard stimuli (10 Hz), windowed within a bar aperture that swept across the visual field in discrete steps. The area outside the stimulus was set to a uniform gray, equal in luminance to the mean of the black and white checkerboards. Both MRI and MEG stimuli were confined to a circular aperture 10° in radius, contrast-reversal rate (10 Hz), bar width (2.5°, *i.e*., 1/4^th^ of the full-field stimulus radius, 10°), but differed in presentation length and sequence (see Experimental design). Details on the stimulus display and experimental design for the MRI and MEG sessions are separately described in the following paragraphs.

#### 3.2.1 Stimulus display - MRI

All subject’s structural and functional data were acquired at the Center for Brain Imaging at New York University. We used a Siemens Allegra 3T head-only scanner for subjects S1 and S2, and a Siemens Prisma 3T full-body scanner for subjects S3-S10 after the NYU Center for Brain Imaging acquired a new scanner. Visual display setup was therefore also different for subjects S1 and S2, compared to subjects S3-S10.

##### Siemens Allegra 3T

For subjects S1 and S2, stimuli were presented with an LCD projector (Eiki LC_XG250, CA, US) with a screen resolution of 1024 × 768 pixels and refresh rate of 60 Hz. Stimuli were displayed onto a translucent back-projection screen in the bore of the magnet. Subjects viewed the screen through an angled mirror mounted onto the coil of the scanner at a distance of ∼58 cm. The stimulus was confined to a circular aperture with a diameter of 20°. The display was calibrated and gamma-corrected using a linearized lookup table.

##### Siemens Prisma 3T

For subjects S3-S10, stimuli were presented with a DPL LED PROPixx projector (VPixx, QC, Canada) with a screen resolution of 1920 × 1080 pixels and refresh rate of 60 Hz. Images were displayed on a translucent back-projection screen in the bore of the magnet. Subjects viewed the screen through an angled mirror mounted onto the coil of the scanner at a distance of ∼83.5 cm. To match the stimuli to previous subjects’ scan sessions, we again confined the stimulus to a circular aperture with a diameter of 20°. The display was calibrated and gamma-corrected using a linearized lookup table.

#### 3.2.2 Stimulus display - MEG

Images were presented using an InFocus LP850 projector (Texas Instruments, Warren, NJ) with a resolution of 1024 x 768 pixels and refresh rate of 60 Hz. Images were projected via a mirror onto a front-projection translucent screen at a distance of approximately 42 cm from the subject’s eyes. The display was calibrated with the use of a LS-100 luminance meter (Konica Minolta, Singapore) and gamma-corrected using a linearized lookup table. The stimuli were confined to a circular aperture with a diameter of 20°.

### 3.3 Experimental design

#### 3.3.1 Experimental design - fMRI

Subjects participated in one 1.5-hr MRI session containing 6 functional runs, where each run was 6.1 minutes. For a given run, the bar apertures show contrast-reversing checkerboard stimuli. The checkerboard contrast pattern oscillated with a 5 Hz square wave, meaning 10 reversals per second. The bar aperture swept across the visual field in discrete steps (1.5s per bar position, 31.5s per bar sweep, see Figure 1A) in 8 different bar configurations (4 different orientations: 0°, 45°, 90°, 135°, with two step directions for each orientation). Two step directions are required for fMRI to avoid biased pRF parameter estimates due to the lag of the hemodynamic response function. After the first, third, fifth and seventh bar sweep, there was a 22.5s mean luminance or ‘blank’ period. In addition, each run started and ended with a 12s blank period. A fixation dot was presented in the center of the screen throughout the run, switching between red and green colors (32 switches per run, average of 7.2s). Subjects were instructed to fixate on the dot throughout the run and report a switch in color with a button press.

**Figure 1.**
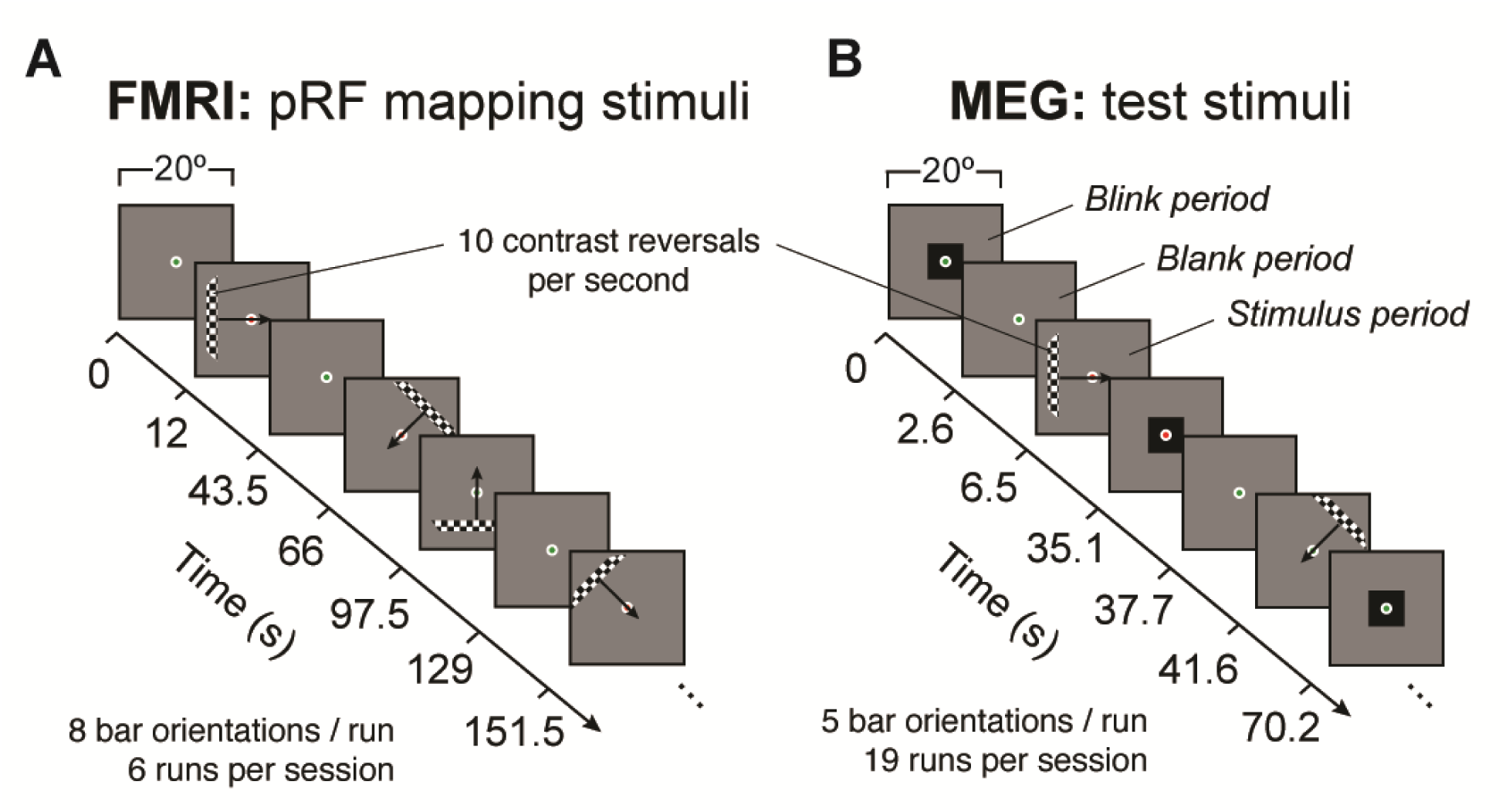
Retinotopic mapping stimuli for fMRI and MEG experiments. **(A)** FMRI stimuli were used to map pRFs on the cortical surface. Contrast-reversing (100% contrast) checkerboard bars swept in discrete steps across the visual field (diameter = 20°, 1 bar step per TR, TR=1.5 s), interleaved with blank periods (mean luminance). One run consisted of 8 bar sweeps along cardinal and off-cardinal axes in both directions. Subjects were instructed to fixate in the center of the screen and press a button every time the fixation dot changed color. Fixation dot is enlarged in this figure for visibility purposes. **(B)** Stimuli presented in the MEG experiment are used in the forward model to create predictions (hence ‘test’ stimuli). Stimuli were similar to fMRI (identical contrast, size, and contrast-reversal rate), except for its sequence and duration. One run contained 5 bar sweeps (3 cardinal, 2 off-cardinal) with shorter bar step durations (1.3 s). Stimulus sweeps were interleaved with blank and blink periods. During blink periods, subjects were encouraged to make eye blinks to limit blinks during blank and stimulus periods. Blink periods were excluded in both data analysis and model predictions.

#### 3.3.2 Experimental design - MEG

All subjects participated in one 2-hr MEG session containing 19 runs, where each run was 3 minutes with short breaks of ∼1 minute between runs. The breaks were terminated when the subject indicted by button press that they were ready for the next run. For a given run, the bar apertures showing the contrast-reversing checkerboard stimuli (10 Hz reversal rate) swept across the visual field in discrete steps (1.3s per bar position, 28.6 s per bar sweep) in 5 different bar configurations for a given run (4 different orientations: 0°, 45°, 90°, 135° with two step directions for 0° and one step direction for 45°, 90° and 135°) (see Figure 1B). MEG runs did not require bar sweeps in both directions for each orientation, because the measured magnetic flux does not contain a time-lag. For the same reason, a randomized sequence might be effective for MEG measurements. Nonetheless, we chose to preserve many stimulus properties matched to the fMRI experiment, while also shortening the experiment to allow for as many repeated runs per subject as possible. As a compromise between shortening and fidelity to the fMRI design, we reduced the number of sweeps from 8 to 5, keeping bidirectional sweeps for one orientation only.

Before every bar sweep and after the last bar sweep, there was a 2.6-s ‘blink’ period indicated by a mean luminance display with a small black square in the center of the screen and then a 3.9-s mean luminance or ‘blank’ period. A fixation dot was presented in the center of the screen throughout the run, switching between red and green colors (32 switches per run, average of 5.6s).

Subjects were instructed to fixate on the dot throughout the run and report a switch in color (every few seconds) with a button press. Subjects were encouraged to blink during the blink period and minimize their blinking during the rest of the run. Blink periods were excluded from analyses.

### 3.4 Data acquisition

#### 3.4.1 Data acquisition - MRI

##### Siemens Allegra 3T

Functional data were collected with a Nova Medical phased array, 8-channel receive surface coil (NMSC072). BOLD fMRI data were acquired using a T2*-sensitive echo planar imaging (EPI) pulse sequence (1500 ms TR, 30 ms TE, and 72° flip angle; 2.5 mm^3^ isotropic voxels, with 24 slices). The slice prescription was placed approximately perpendicular to the calcarine sulcus and covered most of the occipital lobe, and the posterior part of both the temporal and parietal lobes. An additional field map was collected in the middle of the MRI session to correct functional data for B0 field inhomogeneity during offline image reconstruction using an in-house Center for Brain Imaging algorithm.

Structural data were collected in the same (subject S2) or separate MRI session (subject S1) with a Nova Medical head transmit/receive coil (NM011). Data consisted of T1 weighted whole brain anatomical images using a 3D rapid gradient echo sequence (MPRAGE, 1 mm^3^ isotropic voxels). Additionally, a T1 weighted “inplane” image was collected with the same coil and slice prescription as the functional scans to aid alignment of the functional images to the high-resolution T1 weighted anatomical images. This scan had a resolution of 1.25 × 1.25 mm and a slice thickness of 2.5 mm.

##### Siemens Prisma 3T

Both structural and functional data were collected with a 64-channel phased array receive coil. BOLD fMRI data were acquired using a T2*-sensitive echo planar imaging pulse sequence (1-s TR; 30 ms echo time; 75° flip angle; 2 mm^3^ isotropic voxels, multiband acceleration 6). Two additional scans were collected with reversed phase-encoded blips, resulting in spatial distortions in opposite directions. These scans were used to estimate the spatial distortions in the EPI runs and used to correct the EPI runs during preprocessing. Structural data were collected in the same session consisting of T1-weighted whole brain anatomical images (1 mm^3^ isotropic voxels) using a 3D rapid gradient echo sequence (MPRAGE). No additional inplane image was needed for alignment for sessions in the Prisma scanner, because the spatial resolution of the EPIs and the whole-brain coverage enabled direct alignment between the functional images the whole brain T1w anatomical image.

#### 3.4.2 Data acquisition - MEG

MEG data were acquired continuously with a whole head Yokogawa MEG system (Kanazawa Institute of Technology, Japan) containing 157 axial gradiometer sensors to measure brain activity and 3 orthogonally oriented reference magnetometers located in the dewar but facing away from the brain, used to measure environmental noise. The magnetic fields were sampled at 1000 Hz and were actively filtered during acquisition between 1 Hz (high pass) and 500 Hz (low pass).

Before recording, each subject’s head shape was digitized with a handheld FastSCAN laser scanner (Polhemus, VT, USA). Digital markers were placed on the forehead, nasion, left and right tragus and peri-auricular points. To calibrate the digital head shape with the MEG sensor space, five electrodes were placed on the identical location of five digital markers (3 forehead and left/right peri-auricular points). Before and after the main MEG experiment, separate recordings were made of the marker locations within the MEG dewar.

### 3.5 Data analyses

#### 3.5.1 Reproducible computation and code sharing

Nearly all analyses were conducted in MATLAB (MathWorks, MA, USA), except for preprocessing steps converting and preprocessing functional scans from the Prisma MRI scanner using Python. The preprocessing analysis code, MEG forward model and data are publicly available via the Open Science Framework upon publication (URL: https://osf.io/c3hxj/). The code will include scripts to reproduce all figures from the minimally pre-processed data. Each data figure has a single script named *makeFigureX* (where ‘*X’* is the figure number).

#### 3.5.2 MRI Preprocessing

##### Structural data (both MRI scanners)

Structural T1-weighted scans were auto-segmented with FreeSurfer’s recon-all algorithm (Dale, Fischl, & Sereno, 1999; Fischl, Sereno, & Dale, 1999; Fischl & Dale, 2000; Fischl, Liu, & Dale, 2001), available at http://surfer.nmr.mgh.harvard.edu/. For three subjects, small errors in white/gray matter voxel segmentation around the occipital pole were manually corrected. Visually responsive regions of interest (ROIs) were defined on the inflated cortical surface of individual subjects using the probabilistic atlas of visual areas by Wang *et al*. (2015) resulting in boundaries for areas V1-V3, hV4, V3A/B, VO1/2, LO1/2, TO1/2, PHC1/2, IPS0-5, SPL1, and FEF.

##### Siemens Allegra 3T functional data

Using the Vistasoft toolbox available at https://github.com/vistalab/Vistasoft, functional scans were re-oriented to a standardized NIfTI orientation (RIA to LAS), slice-time corrected by resampling the time series in each slice within the 1.5s-volume to the center slice, and motion corrected by aligning all volumes of all scans to the first volume of the first scan using 3D rigid body alignment (6 DOF). The first 8 volumes of each functional scan were removed to avoid unstable magnetization of the scanner. Functional scans were aligned to the T1-weighted anatomical scan using a coarse, followed by a fine, 3D rigid body alignment with the additional inplane scan (Vistasoft’s *alignvolumedata_auto*).

##### Siemens Prisma 3T functional data

Functional scans were converted from dicom into BIDS format (Gorgolewski et al., 2016) using NYU Center for Brain Imaging in-house version of NIPY’s heudiconv, available at http://as.nyu.edu/cbi/resources/Software.html. The following in house preprocessing workflow was implemented with the nipype toolbox (Gorgolewski et al., 2011), and is available via GitHub (https://github.com/WinawerLab/MRI_tools/blob/master/preprocessing/prisma_preproc.py). Using the FSL toolbox (Smith et al., 2004), all volumes from all EPIs were realigned to the single-band reference image of the first EPI scan. This single band reference image was then registered to the additional spatial distortion scan with the same phase encoding direction. The two additional spatial distortion scans with opposite phase-encoding direction were then used to estimate the susceptibility-induced warp field using a method similar to (Andersson, Skare, & Ashburner, 2003). Motion correction (3D rigid body, 6 DOF), registration to the spatial distortion scan and unwarping were then applied in a single step to each volume of each EPI. The unwarped EPIs were aligned to the high-resolution whole-brain T1 using FreeSurfer’s *bbregister* (6 DOF, rigid).

##### Siemens Allegra & Prisma 3T functional data

Time series from EPIs were resampled to 1 mm^3^ isotropic voxels, *i.e.,* the resolution of T1-w anatomy, within the gray matter voxels using trilinear interpolation. This step allows for easy comparison of functional to anatomical data using FreeSurfer’s tools. Time series within the gray matter voxels were converted into percent signal change by dividing the signal by its mean. Baseline drifts were removed from each run with high-pass temporal filtering using 3 discrete cosine terms (0 cycles or ‘DC’; 0.5 cycle and 1 cycle). At last, all 6 runs were averaged given that subjects saw the same stimuli within a dataset.

#### 3.5.3 MEG Preprocessing

The FieldTrip toolbox (Oostenveld, Fries, Maris, & Schoffelen, 2011) was used to read the raw data files. For all subsequent MEG analyses, custom code was written in MATLAB. With use of the triggers from the stimulus presentation computer, MEG data were first divided into 1300 ms epochs (*i.e.,* matching the duration of 1 bar step) for every MEG sensor. For all subjects, epoching resulted in an initial 2660 epochs per sensor: 22 consecutive epochs per bar sweep, with 2 consecutive epochs for blink and 3 consecutive epochs for blank periods before each sweep, and after the last bar sweep of every run, 5 bar sweep directions, for 19 runs. To avoid the transient response associated with a change in the stimulus (either a change in bar position or from a blank period to a stimulus period), we then shortened each epoch to 1100 ms, skipping the first 150 ms and last 50 ms of each 1300-ms epoch. We choose to remove the first 150 ms to skip one full cycle of the 10 Hz response (100 ms) plus 50 ms to approximate the time for the neural response to reach the cortex. The last 50 ms were removed so that the total epoch length was an integer number of cycles at the steady-state frequency (10 Hz).

Outlier epochs were removed in the following way. First, epoched data were high-pass filtered with a 1 Hz Butterworth filter (with a high-pass amplitude of 3 dB and a passband frequency of 0.1 Hz and amplitude of 60 dB). We then computed the variance within every 1100-ms epochs (across time points), for each MEG sensor. We labeled an epoch as ‘bad’ if its variance was 20 times smaller or 20 times larger than the median variance across all epochs and sensors. If more than 20% of the epochs were labeled bad for a given sensor, then we removed the entire sensor from analysis. If more than 20% of sensors contained the same ‘bad’ epoch, we removed the entire epoch from analysis (*i.e.,* for all sensors). These criteria succeed in identifying known outliers (5 sensors that had long-term hardware problems as well as sensor/epoch combinations in which the responses became temporarily saturated due to external noise), while at the same time avoiding the removal of unnecessarily large amounts of data. If a given epoch was labeled ‘bad’ for some sensors, but was not removed for all sensors, the data of the removed sensors were replaced by the time series spatially interpolated across nearby sensors (weighting sensors inversely with the distance). We removed on average ∼21% of each dataset, including all epochs from the 5 sensors with long-term hardware problems. We tested the effects of changing the variance thresholds for removing individual sensor/epoch combinations, and the effects of changing the criteria for removing entire epochs (all sensors) or entire sensors (all epochs) in an example subject with intermediate data quality (S5). Using the same settings as for all other subjects resulted in 13.9% of data being labeled as ‘bad’, including 6 sensors. Adjusting the lower variance bound did not affect the percentage of data labeled as ‘bad’. For the upper variance bound, a more liberal (10x) or more conservative (40x) threshold either increased by 7.6% or decreased by 1.6% percentage of ‘bad’ labeled data respectively. Increasing the percentage to mark entire sensors or epochs as ‘bad’ did not affect the number of ‘bad’ sensors and a less than 1% decrease in ‘bad’ epochs. Decreasing the percentage from 20 to 10% (so more liberal) marked an additional 2 sensors and ∼3% of epochs as ‘bad’. Importantly, none of these changes in outlier criteria caused a substantial change in model performance nor affected our results.

We used the Noisepool-PCA algorithm to increase the signal-to-noise ratio (SNR) of our MEG time-series (Kupers et al., 2018). This algorithm was adapted from an fMRI algorithm called GLMdenoise (Kay, Rokem, Winawer, Dougherty, & Wandell, 2013). In short, for each subject the algorithm defines a noise pool: a subset of sensors that contains little to no 10 Hz visually evoked steady-state response. Time series within each epoch and sensor of the noisepool were then filtered to remove all 10 Hz (and harmonics) components. Using principal components analysis (PCA), we defined global noise regressors from the filtered noise pool time series. For each subject, the first 10 PCs were used to create 10 new denoised datasets: the first denoised dataset had the PC 1 projected out of the data in each sensor, epoch by epoch. The second denoised dataset had PC1 and PC2 projected out, etc. For each denoised dataset, we calculated the median R^2^ across bootstrapped epochs. The optimal number of PCs to project out was the smallest number of PCs that resulted in a denoised data with a median R^2^ within 5% of the maximum possible median R^2^ of 10 datasets. This resulted in removing 6 PCs on average across subjects, ranging between 2-8 PCs. At last, we reshaped the denoised MEG data into a 4D array: *t* (time points) × *k* (epochs) × *m* (sensors) × *r* (runs).

#### 3.5.4 MEG data quality check

We calculated two parameters to check the quality of the measured MEG data: 10 Hz coherence and split-half reliability of the 10 Hz steady-state visually evoked responses. The coherence of the 10 Hz steady-state visually evoked fields (SSVEFs) provides an estimate of the signal-to-noise ratio of the steady-state response within stimulus periods. The 10 Hz SSVEF coherence was defined by dividing the average 10 Hz amplitude across epochs of all runs by the average amplitudes of 10 Hz and neighboring frequencies (*i.e.,* 9 to 11 Hz) across epochs of all runs.

The second metric was the split-half reliability of the 10 Hz steady-state amplitudes, providing an estimate of how reliable the steady-state responses are across runs. We computed the split-half reliability by dividing the 19 repeated runs into two groups. After taking the sensor-wise average time series across runs for each of the two data splits, we applied the FFT to the two run averages and extracted the 10 Hz amplitude per epoch. The 10 Hz amplitudes for the first data half were then pairwise correlated to the 10 Hz amplitudes for the other data half (Pearson’s ρ). This split-half reliability procedure is repeated 1000 times and summarized as the mean correlation across repetitions, resulting in one split-half reliability sensor map per subject.

#### 3.5.5 MRI-MEG head model and alignment

The head model, also referred to as the ‘lead field’ or ‘gain matrix’, describes the contribution of cortical locations (or ‘sources’) to the activity at each individual MEG sensor. To generate this head model, we align the individual’s anatomy and the MEG helmet in a common coordinate space using the Brainstorm toolbox (Tadel, Baillet, Mosher, Pantazis, & Leahy, 2011).

Specifically, we defined the nasion and left/right peri-auricular points in the T1-weighted image of each individual subject. We used Brainstorm’s automated alignment algorithm to align the fiducials marked in the T1-weighted image, the recorded locations of electrodes attached to the subject’s face while lying in the MEG scanner, and points in the 3D head shape. Small manual translational adjustments were applied to the rotation matrix if necessary. After alignment, we computed the individual subject’s head model using Brainstorm’s implementation of the overlapping spheres algorithm (Huang, Mosher, & Leahy, 1999) using the subject’s FreeSurfer pial surface (∼300,000 vertices per hemisphere). The overlapping spheres algorithm fits a different sphere to the subject’s skull for each sensor. We choose the overlapping spheres algorithm for its low computational cost while having an accuracy comparable to the more biologically accurate but computationally demanding Boundary Element Model (BEM) (Kybic et al., 2005; Gramfort, Papadopoulo, Olivi, & Clerc, 2010). We did not downsample the number of vertices as is often a standard implementation in MEG/EEG software packages, as we do not need to reduce computational cost for our forward model (in contrast to inverse modeling), enabling us to avoid interpolation errors introduced by downsampling of the pRF parameters from a high to a low-resolution cortical surface. We constrained our head model to one perpendicular dipole per vertex, resulting in a matrix of FreeSurfer vertices (∼300,000, depending on the subject) by 157 sensors.

### 3.6 A stimulus-to-sensor model for MEG responses

We developed a modelling framework that learns cortical pRFs from fMRI data, and then uses a biophysics model (gain matrix) from anatomical MRI co-registered to MEG data. The model takes as input a visual stimulus and predicts as output the MEG SSVEF amplitude at each sensor and each stimulus position. The voxel-wise pRF parameters, fit to fMRI data, are projected to the cortical surface and used to predict neural population responses to the MEG stimuli. These predicted values are in the form of one number per voxel per bar position. Because both our stimulus and our pRFs are defined as non-negative, the predicted cortical responses are all also non-negative. These predictions are then projected to the MEG sensor space using the gain matrix from the overlapping-spheres head model (Huang et al., 1999). The values projected to the sensors are signed because the gain matrix is signed. These predicted MEG data are compared to the measured phase-referenced steady-state MEG response using a linear regression, fitting a reference phase *θ*_ref_ and a gain parameter *β* per sensor to maximize the coefficient of determination (R^2^) (Figure 2, training model). The optimal reference phases were then cross-validated across data halves to recompute the phase-referenced 10 Hz steady-state responses and averaged across halves. The corresponding gain factors were averaged across halves and used to scale the initial predicted sensor responses. A final goodness of fit of the average predicted MEG responses was computed on the average measured MEG responses (Figure 2, test model). We explain each of these steps in detail below.

**Figure 2.**
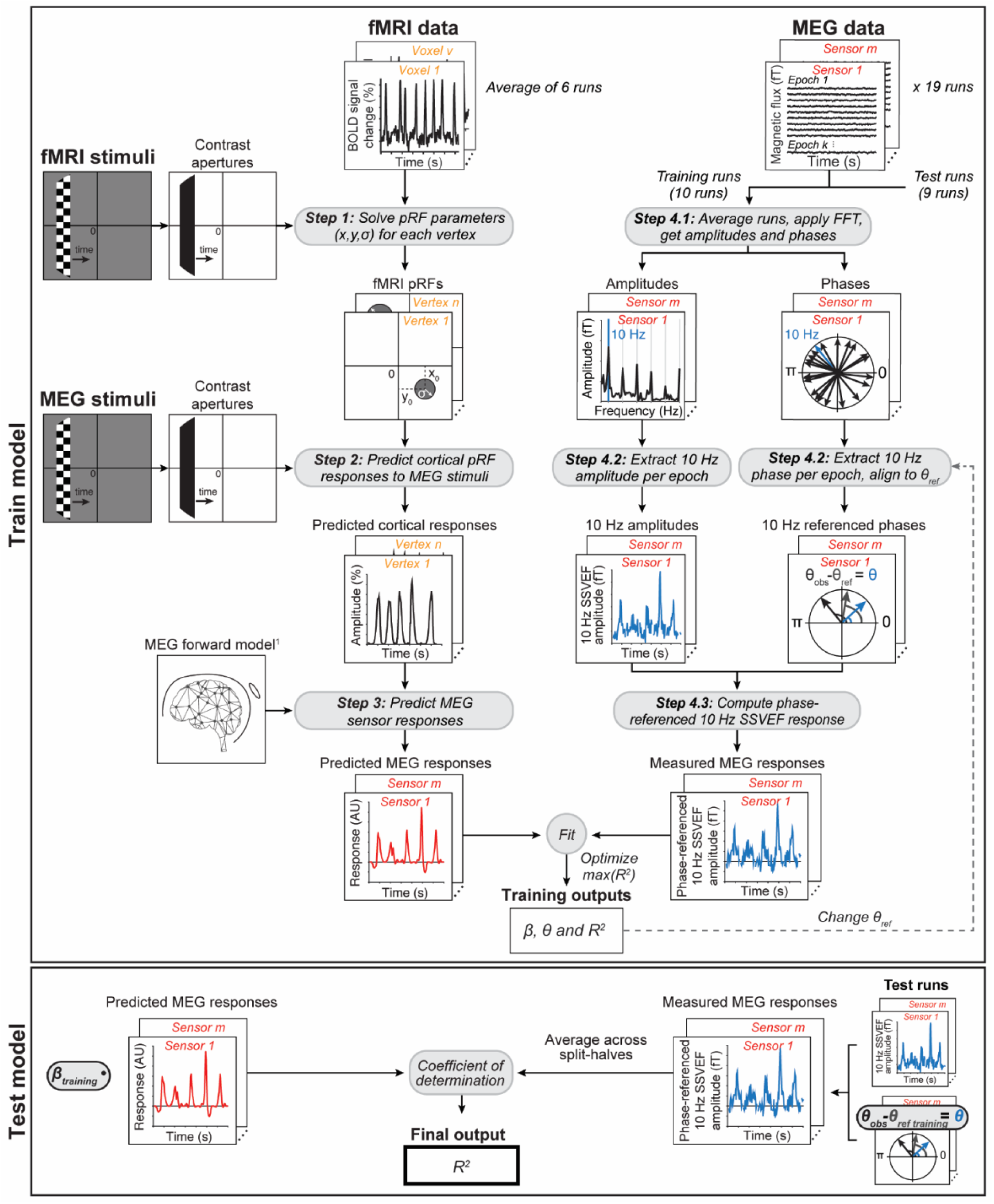
MEG forward modeling approach. The model starts with preprocessed fMRI and MEG data and their corresponding stimuli as inputs. **Train model. Step 1:** FMRI stimuli are binarized into apertures and used to solve pRFs within each cortical location and projected to the cortical surface. **Step 2:** Estimated pRFs are multiplied with MEG stimulus apertures to predict time series on the cortical surface. **Step 3:** Predicted cortical responses are multiplied with the gain matrix from the MEG forward model to get predicted MEG responses. The gain matrix describes the contribution of each source to magnetic fields measured in MEG sensors and is computed by the overlapping spheres algorithm (^1^Huang *et al*., (1999)). Predicted responses are fitted to measured MEG responses, using a split-half cross-validation procedure. **Step 4.1:** MEG training runs are averaged and its time series are transformed to the Fourier domain. **Step 4.2:** 10 Hz amplitudes and phases are extracted per epoch and sensor. **Step 4.3:** 10 Hz phase and amplitudes are combined into phase-referenced 10 Hz SSVEF amplitudes by fitting the predicted MEG responses from pRFs to measured MEG responses. This model fit uses two free parameters (gain *β* and reference phase *θ_ref_*) and is optimized by finding the reference phase where the prediction explains most variance in the data. **Test model.** Both free parameters are cross-validated: the optimal reference phases from training are used to compute phase-referenced 10 Hz SSVEF responses of the test runs as in Step 4. The gain parameters are summarized by the weighted average across the two training iterations and used to scale the predicted MEG responses. At last, measured MEG responses are averaged across split-halves and compared to predicted MEG responses using the coefficient of determination (R^2^).

#### 3.6.1 Step 1.1: Solve pRFs with fMRI

Using the Vistasoft toolbox (https://github.com/vistalab/Vistasoft), we solved linear, circularly symmetric 2D Gaussian pRF models on the functional MRI data, as previously described in Dumoulin and Wandell (2008). In brief, pRF models were solved by a two-stage coarse-to-fine optimization procedure on the gray matter voxels, using the binarized MRI stimulus apertures and Vistasoft’s built-in ‘difference between two gammas’ hemodynamic response function. The first stage of the optimization procedure started with a coarse grid-fit. The best fitting parameters for each voxel from the coarse grid-fit were used as the seed for the fine grid-fit. This fitting procedure resulted in an estimated preferred pRF size (*σ*, 1 SD of 2D Gaussian), center location (*x*, *y*), gain (or scaling factor), and variance explained for every voxel. The pRF parameters computed at gray matter voxels are interpolated to surface vertices.

#### 3.6.2 Step 1.2: Smooth pRF parameters across gray matter voxels

The pRF parameters are interpolated from the gray matter voxels (*i.e.*, voxels comprising the ‘cortical ribbon’) to the surface vertices using a nearest neighbor interpolation algorithm. This choice was made because of technical constraints within the Vistasoft toolbox. One could alternatively change the order of operations and first interpolate the time series to the surface and then solve the pRF parameters. The results would likely be similar in that we used nearest neighbor interpolation to project pRF solutions from cortical voxels to surface nodes. To reduce sensitivity to noise, we smooth pRF parameters across the cortical surface by calculating a weighted average over a normalized truncated gaussian kernel (Andrade et al., 2001). This procedure applies surface-oriented smoothing using the geodesic rather than Euclidean distance, respecting the topology of the cortical surface. The Gaussian kernel (approximately, a FWHM of 3 mm at 1 cubic mm of voxel resolution) is created at every gray matter vertex. Neighboring vertices in which estimated pRF model fit did not explain any variance of the data (*i.e.,* a variance explained of 0%) were excluded. We smoothed the position (*x*, *y*) and size (*σ*) parameters as well as a proxy for the pRF gain (scale factor, or “beta weight” in the Vistasoft code). Although the pRF model is linear, it is not linear with respect to its parameters, and smoothing of the parameters can have unwanted effects, particularly in the amplitude of the response (controlled by the pRF gain). This is due in part to the fact that the software implementation defines the pRFs as Gaussians with unit height at the pRF center, such that the pRF volume within the stimulus aperture depends on the size of the pRF and the degree of overlap between the pRF and the stimulus aperture. To ensure that the smoothing procedure resulted in smoothing of the time series amplitudes, we used the pRF models to predict the time-course amplitude (using the fitted beta parameter), and then smoothed the maxima of these predicted time-course amplitudes over the surface.

#### 3.6.3 Step 2: Predict neural responses to MEG stimuli from pRF parameters

To predict the steady-state responses in MEG sensors, we first created a predicted response from estimated pRF parameters on the cortical vertices. Vertices were constrained by those whose pRF parameters explained more than 10% of the variance in the MRI data. The 10% threshold was chosen to exclude vertices that are likely to reflect noise and are not visually responsive (or incoherent with the stimulus). Moreover, we restricted the vertices to only those whose pRF centers fell within our stimulus aperture (10° of visual angle), and those which fell inside the visual ROIs from Wang *et al*.’s probabilistic atlas (2015). For all other vertices, the predicted response was set to 0. For each vertex, a 2D Gaussian receptive field was constructed using its preferred center and size. The height of this receptive field was scaled by the vertex’s pRF gain. A dot product of these receptive fields and the binarized MEG stimulus resulted in the predicted surface response (one value per aperture position). As mentioned earlier, blink periods were excluded. Blank periods were predicted as zero responses, assuming that blank screen epochs elicit a negligible 10 Hz steady-state visually evoked response with a random phase. Vertices with a maximum predicted response larger than 10 times the median of all vertex maximum responses were considered outliers and excluded. This criterion was implemented ad hoc, after investigating the predicted time series of individual subjects and finding several pRF time series with unrealistically large amplitudes (>100% signal change). This threshold had no effect on two datasets and removed a very small amount of data for the other eight datasets (less than 0.3% of vertices with a predicted cortical response per subject).

#### 3.6.4 Step 3: Predict MEG sensor responses from neural responses

The matrix containing the predicted pRF responses on the cortical surface S was then multiplied with the gain matrix from the MEG head model G, resulting in predicted MEG sensor responses Ŷ (**Equation 1**). We compared these predicted MEG sensor responses to the measured MEG responses.

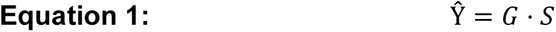

Where Ŷ *(k* epochs × *m* sensors) are the predicted pRF responses for MEG sensors, G (*n* vertices × *m* sensors) is the head model, and S (*k* epochs × *n* vertices) is the predicted pRF response on the cortical surface.

#### 3.6.5 Step 4: Fitting the model to MEG training data

The observed MEG responses were computed as the phase-referenced 10 Hz steady-state visually evoked fields (SSVEFs), using cross-validation. Phase referencing the amplitude is done when the expected oscillations can be either positive or negative, which can occur because the gain matrix created by the head model is signed (*i.e.,* contains both positive and negative numbers). Moreover, the reference phase itself may be of interest, as it can capture differences in timing between sensors driven by different regions of cortex, with different response properties.

For each subject, epoched MEG data were split into two halves: a training half containing the 10 odd runs and a test half containing the 9 even runs. We then computed the sensor-wise average time series within each epoch across training runs and transformed the average to the Fourier domain by applying the FFT to the time series data (Figure 4, Step 4.1). We extracted both amplitude and phase information from the spectral MEG data at 10 Hz (*i.e.,* the contrast-reversal rate of the stimulus) to compute a *phase-referenced steady-state response* (Figure 4, Step 4.2). To calculate this response, we describe the 10 Hz Fourier component of a given epoch as a vector with amplitude length and phase angle (*i.e.,* cosine of the phase). We then scale the 10 Hz amplitude by the difference in angle between the measured phase and a reference phase *θ*_ref_ resulting in the phase-referenced steady-state responses Y, for every sensor *m* and epoch *k* (Figure 4, Step 4.3). The reference phase *θ*_ref_ was obtained separately for every sensor by choosing the phase leading to the highest variance explained in the MEG time series after iterating over all 100 possible reference phases. The variance explained was computed by a linear regression of the model predictions to the phase referenced time series with one free parameter *β* (*i.e.,* a scale factor or gain). This scale factor brings the predicted time series into units of femto-Tesla. Fits were optimized by maximizing the coefficient of determination (R^2^) between model and data (*i.e.,* the residuals sum of squares divided by the total sum of squares). After iterating over all possible reference phases, we choose the one whose MEG time series was best matched to the predicted MEG responses by linear regression constrained to positive scale factors (largest R^2^). (We constrain to positive scale factors for consistency, because for every reference phase, there is another phase 180° apart which makes the identical predictions up to a sign flip).

#### 3.6.6 Test model: Comparing predicted to measured MEG responses

The model predictions were tested using a split-half cross-validation approach. Once the optimal reference phases were selected for every sensor for the *training* half, they were applied to compute the phase-referenced MEG 10 Hz steady-state response in the *test* half (see **Equation 2**). This was repeated for each of the two split halves.

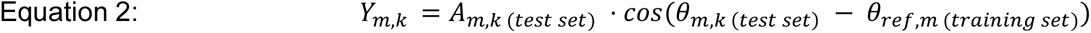

Where Y*_m,k_* is the phase-referenced MEG 10 Hz steady-state response of the test runs, for every epoch *k* and sensor *m*. A*_m,k_* is the average 10 Hz amplitude across test runs, for every epoch *k* and sensor *m. θ_m,k_* is the average 10 Hz phase across all test runs, for every epoch *k* and sensor *m*. *θ_ref_*_,*m*_ is the reference phase for every sensor *m*, computed by fitting the training data to the predicted responses.

Training both data halves resulted in two sets of *β* parameters corresponding to the model fit with the optimal reference phase. Because the predicted cortical responses were identical for both data halves, we scaled the predicted MEG responses with the weighted average of the two *β* parameters, resulting in one predicted time series per sensor. We used a weighted average because the two halves had an unequal number of runs.

The entire cross-validated phase-referencing procedure resulted in two arrays: one with phase-referenced SSVEF responses (*k* epochs × *2* groups of runs × *m* sensors) and one with scaled predicted MEG responses (*k* epochs × *m* sensors). Measured MEG data were averaged across the two run groups, resulting in a matrix of *k* epochs × *m* sensors. To summarize the goodness of fit across the entire data set, we computed the coefficient of determination (R^2^) for the average predicted MEG responses to the average measured MEG responses.

#### 3.6.7 Sensitivity of prediction to rotating pRF centers

To evaluate how sensitive our model predictions are to pRF parameters estimated by fMRI, we systematically altered the fMRI pRF parameters. We estimate the sensitivity to pRF center position by systematically rotating the pRF centers around the fovea. We do so by first calculating the polar angle for a given vertex using the *x* and *y* pRF parameters, and then adding an angle rotation from –180° to 180° in one of 8 equal steps of 45°. For every rotation, we fit and test the model in the same way as we did for the data without rotation, including fitting the reference phase and scale factor per sensor and evaluating by cross-validation.

#### 3.6.8 Sensitivity of prediction to scaling pRF sizes

We estimate the sensitivity of our model to pRF size by systematically scaling the originally estimated pRF size (*σ*). We scaled original pRF sizes from 5 times smaller to 10 times larger, in 19 log-spaced steps, where a scale factor of 1 is the pRF size estimated with fMRI. Similar to the rotation manipulation, we re-computed the predicted MEG responses and optimal reference-phases after applying a particular scale factor.

#### 3.6.9 Sensor selection for summarizing results pRF position or size manipulations

To evaluate the effects of rotating pRF centers and scaling pRF size, we average the variance explained across a subset of sensors for each subject. We use two approaches for sensor selection to check for robustness of our results. One approach is to use model accuracy. To select the subset of sensors, we take the union of the 10 sensors with highest variance explained by the model in each of the rotation steps or each scaling step. This results in a minimum of 10 sensors per subject, but typically more because the top 10 sensors are not the same across the pRF manipulations. This selection method is unbiased toward any particular rotation angle or scale factor and is agnostic about the spatial location of the sensors. By selecting the top 10 sensors, we avoid including large amounts of noise from visually unresponsive sensors. We chose 10 out of consistency with prior work (Kupers et al., 2018) and because it approximately matched our visual inspection of the number of responsive sensors. We also checked the effects of selecting the top 5 or the top 15.

The second approach is data-driven and does not use the pRF model. For this approach, we select the 10 sensors with the highest split-half reliability of the 10 Hz SSVEF signal. This results in exactly 10 sensors per subject.

Data were summarized for individuals by selecting sensors using one of the two approaches described above. This resulted in a matrix of variance explained (number of rotation angles or scale factors by selected sensors). We then take the mean and standard error of the mean across selected sensors for each rotation angle or scale factor as our summary metric.

### 3.7 Group average model fits

A challenge in group analysis of MEG or EEG is that the same sensor in two subjects do not sample from the identical parts of the brain. An advantage of a forward model of the MEG signal is that group average data can be computed in the model space, fit separately for each subject. The sensor-wise average prediction across subjects accounts for the differences in cortical sampling between subjects, because each prediction is based on that subject’s fMRI data, head model, and MEG training data. The average prediction can then be compared to the average group data. We refer to this method as **average-then-goodness-of-fit**. This method provides a compact summary of the results in sensor space. However, the interpretation is not straightforward since the same sensors do not pool from the same brain regions across subjects.

We computed the **average-then-goodness-of-fit** group result by taking each subject’s cross-validated predicted MEG responses (thus scaled by the individual subject’s gain factors, *β*) and observed MEG responses (phase-referenced using a reference phase *θ*_ref_ optimized to the individual subject’s predicted MEG responses). We then average the predicted MEG responses across subjects and separately average the observed MEG responses across subjects, resulting in two matrices: both *k* epochs by *m* sensors. We compare the goodness of fit using the coefficient of determination. In the case where we altered the pRF parameters, for each rotation or scaling iteration, we bootstrapped the average measured and predicted MEG responses across subjects 10,000 times. We compute the coefficient of determination between the two averages for each bootstrap, resulting in a variance explained distribution for each sensor. From this distribution, we extracted the mean variance explained and the 14^th^ and 86^th^ percentile for upper and lower bounds of the 68%-confidence intervals.

We also implemented a second group average which reverses the order of operations. Rather than computing the model accuracy of the averaged data, we compute the average model accuracy across individual subjects. We refer to this method of computing the group average as **goodness-of**-**fit-then-average**. In contrast to the first method, the sensors used to compute model accuracy for this method differ across subjects. Again, we bootstrapped across subjects 10,000 times and computed the average and 68%-confidence intervals across bootstraps.

#### 3.7.1 Model predictions from group average pRF parameter maps

The forward model could also be implemented without collecting subject specific fMRI, for example with a retinotopy template or average group data from a different study. For comparison, we derived average pRF parameter maps from an aggregate 3T retinotopy dataset with 44 subjects (Himmelberg, Kurzawski, *et al*. (2021)). Data were collected at the same NYU scanner facility using the 3T Prisma MRI scanner with approximately the same field-of-view as the MEG experiment, but with different stimuli. Subjects in this aggregate dataset viewed 6 runs of colorful sweeping bar stimuli, similar to those used for the Human Connectome Project 7T retinotopy dataset (Benson et al., 2018). PRF models were solved within individual subjects using the same Vistasoft software. The *x*, *y*, and *σ* parameter maps of each individual subject were interpolated onto a template cortical surface (FreeSurfer’s *fsaverage)* and then averaged across subjects. These group-average parameter maps were then interpolated onto each of our 10 original individual subject’s mid-gray cortical surface using Neuropythy’s *interpolate* with the default nearest-neighbor method (https://github.com/noahbenson/neuropythy) (Benson & Winawer, 2018). Both interpolation steps–the 44 individuals mapped onto *fsaverage* to create the template, and the application of the template to the individuals in the MEG experiment–used nearest-neighbor interpolation.

While it is reasonable to assume that the *x*, *y*, and *σ* pRF parameters will be similar for the large retinotopy dataset and for the MEG experiments, the gain may differ substantially. For example, the fMRI retinotopy dataset was measured with colorful static stimuli containing objects and textures, whereas the stimuli for the MEG experiment were achromatic moving checkerboards. Different visual field maps may be more responsive to one of these stimulus types than the other. For this reason, we did not compute pRF scale factors from the NYU retinotopy database with 44 subjects. For simplicity, we assumed that the response gain was uniform within a map (each ROI in the Wang et al. (2015) atlas) but could differ between maps. To derive a gain for each map, we averaged the response gain across the 10 subjects with fMRI data collected for this paper with drifting checkerboards. We took the median response from voxels within a map for each subject, defining the response as the maximum predicted percent signal change in the predicted cortical time series. This resulted in a matrix of median values, with a size of 10 subjects by 25 ROIs. Median values were then averaged across subjects per ROI. To apply the template in our forward model, all vertices within a given ROI were given the corresponding average gain value. This cortical map was used to scale the maximum predicted cortical pRF responses reconstructed from the average *x*, *y*, and *σ* pRF parameter maps from the aggregate NYU 3T retinotopy dataset. Once the average pRFs were reconstructed on an individual subject’s cortical surface, all following analysis steps of our forward model were identical.

## 4 Results

In separate MRI and MEG sessions, subjects viewed high contrast retinotopic bar stimuli traversing across the visual field, where the checkerboards inside the aperture reversed contrast 10 times per second. Data from the MRI session were used to reconstruct population receptive fields (pRFs) on the cortical surface for each individual subject, using the modeling approach described by Dumoulin and Wandell (2008). These pRFs on the cortical surface were the building blocks of our forward modeling approach, as they were used to predict the observed MEG response. Below we describe the observed steady-state components within the MEG data and report the MEG forward model performance using the pRFs estimated from the MRI session. Finally, we show the effect of artificially altering the initially estimated pRFs on our MEG model.

### 4.1 Retinotopic stimuli produce reliable steady-state responses in posterior MEG sensors

MEG data from individual subjects were divided into 1.1-s non-overlapping time bins (epochs), for every sensor and run. These epochs contained either a contrast-reversing bar at a particular location in the visual field (‘stimulus periods’), a zero-luminance screen (‘blank periods’), or a square stimulus prompting subjects to rest and make excessive eye blinks (‘blink periods’). The latter were removed from all following analyses. Both stimulus and blank periods were averaged across multiple runs, before transforming the MEG time series to the Fourier domain.

We found a large steady-state response at 10 Hz (the contrast-reversal rate of the stimuli) and multiples of 10 Hz (*i.e.,* harmonics) during stimulus periods compared to blank (Figure 3A). These 10 Hz steady-state visually evoked fields (SSVEFs) were largest in posterior MEG sensors. To estimate how robust 10 Hz steady-state responses were across identical stimulus runs, we computed two data metrics of the 10 Hz amplitudes: its coherence and split-half reliability.

**Figure 3.**
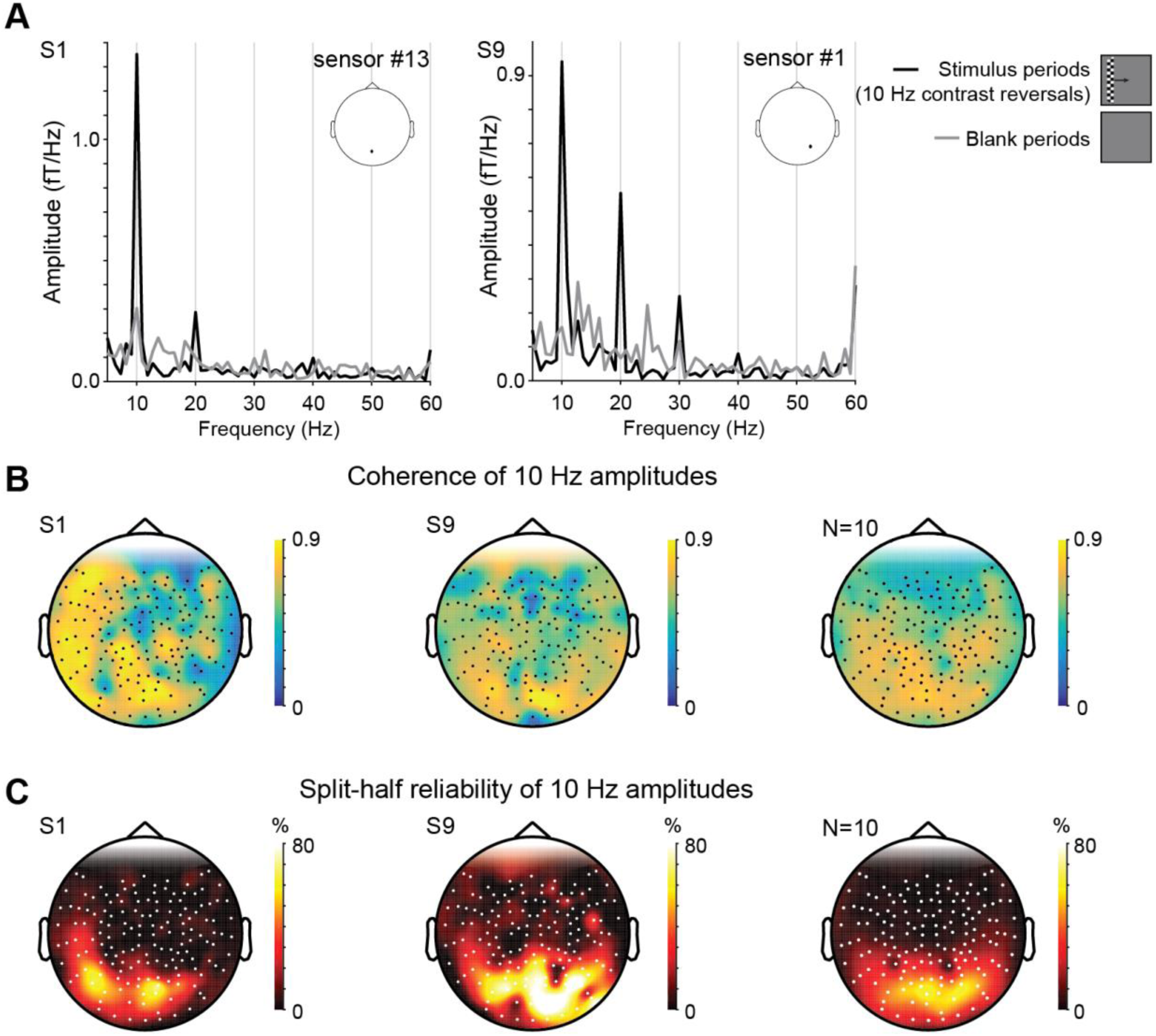
Steady-state visually evoked amplitudes from the MEG experiment. **(A)** Example spectra from two posterior MEG sensors (location indicated by dot on schematic head) and two subjects (S1 and S9). Fourier transformed stimulus periods (black line) show a large peak at the contrast reversal rate (10 Hz, *i.e.,* the steady-state visually evoked field or ‘SSVEF’) and multiples of 10 Hz (harmonics) compared to blank periods (grey line). Note that these amplitudes contain only positive values and are not yet referenced by the corresponding phases. **(B)** MEG sensor topography of 10 Hz SSVEF coherence (10 Hz amplitude divided by mean of 9 to 11 Hz amplitude) for subjects S1 and S9 and sensor-wise average across all subjects (N=10). **(C)** Split-half reliability of the 10 Hz SSVEF amplitudes for subjects S1 and S9 and sensor-wise average across all subjects (N=10).

The coherence metric provides a signal-to-noise ratio of the steady-state response within stimulus periods for every MEG sensor, without regard to the particular stimuli giving rise to the response. This metric is computed by dividing the average 10 Hz amplitude of all stimulus periods by the sum of the amplitudes from 9 to 11 Hz. We found that the coherence of the steady-state response is largest in posterior MEG sensors (Figure 3B), in line with the expectation that posterior sensors are located over the visual cortex and maximally driven by the stimulus contrast-reversals.

The specific 10 Hz coherence sensor topography varied across subjects. For example, subject S1 (Figure 3B, left panel) showed extended regions of high 10 Hz coherence in lateral and anterior MEG sensors, whereas subject S9 did not (Figure 3B, middle panel). When sensor-wise averaging 10 Hz coherence topographies across subjects, the coherence values are highest in posterior sensors (Figure 3B, right panel). This indicates that across subjects 10 Hz steady-state amplitudes are most robust in posterior MEG sensors, as expected due to proximity of these sensors to visual cortex.

To estimate how reliable the 10 Hz steady-state amplitudes are across the 19 repeated runs in the MEG experiment, we computed the split-half reliability. Unlike the coherence metric, which average across epochs, the split-half reliability was sensitive to the specific pattern of responses as a function of bar position. We found that split-half reliability is largest in posterior MEG sensors (up to Pearson’s *ρ* = ∼80%) in both individual maps and across-subjects maps (see Figure 3C). Many posterior sensors with high reliability overlap those sensors with the largest coherence within individual subjects (see Supplementary Figure S1). The sensors with high 10 Hz coherence tend to spread out more to lateral and frontal MEG sensors compared to those with high split-half reliability, which are confined to posterior MEG sensors. A possible explanation for this topography discrepancy is that some sensors in anterior locations are broadly sensitive to the stimulus (high coherence) but have little to no position sensitivity (low split-half reliability).

### 4.2 Forward model predicts phase-referenced MEG responses in posterior sensors

Thus far, we focused on the 10 Hz steady-state spectral *amplitudes* and ignored the corresponding 10 Hz *phases*. This phase component can vary across epochs and MEG sensors due to processing delays in the visual system and depend on stimulus features, such as contrast (Shapley & Victor, 1978) and eccentricity (Jeffreys, 1971; Burkitt, Silberstein, Cadusch, & Wood, 2000; Ales, Yates, & Norcia, 2013; Inverso, Goh, Henriksson, Vanni, & James, 2016). Because our stimulus was a bar sweeping in different directions across the visual field and likely activated both early and later visual areas which differ in response timing, we expected variability in the 10 Hz phases across MEG sensors. Additionally, the gain matrix from the MEG head model is signed, causing predicted MEG responses to be signed. Therefore, to use all available information in the MEG data, we combined 10 Hz amplitudes and phases into 10 Hz phase-referenced steady-state responses. We did so by scaling the 10 Hz amplitudes by the cosine of the difference between the observed phase and a reference phase (see **Methods**). This way both predicted responses and measured MEG responses are signed.

To predict the MEG responses to retinotopic stimuli for each individual subject, we developed a forward model (Figure 2). In short, our forward model predicted the MEG responses for every sensor by first multiplying pRF models estimated from fMRI at every cortical location with the MEG stimulus, for every time point. We then multiplied the resulting pRF time series at every cortical location with the gain matrix from the MEG head model based on subject’s anatomy and head position in the MEG. For the measured MEG responses, we combined amplitude and phase information into a phase-referenced amplitude for every sensor. We used split-half cross-validation to determine the optimal reference phase for every MEG sensor by fitting observed MEG responses to the predicted MEG responses, optimizing for variance explained by the model. By splitting the MEG runs into two groups, reference phases of the first half were used to compute the phase-referenced SSVEFs for the second half. Finally, to determine the overall goodness of fit of the model, we compared the predicted time series with the observed phase-referenced 10 Hz SSVEFs averaged across both split-halves for every MEG sensor.

By combining local pRFs on the cortical surface with the biophysical head model, our forward model was able to capture ∼60% of the variance in phase-referenced steady-state MEG data in posterior MEG sensors (Figure 4). The predicted MEG responses in sensors with high variance explained usually contained five peaks across the 154-s experiment, corresponding to the five orientated bar sweeps across the visual field. This result was found both at the group level (Figure 4B, left panel), as well as individual subject level (Supplementary Figure S2). Those MEG sensors with highest variance explained by the forward model approximately overlap with subset of posterior sensors that contain large 10 Hz coherence and split-half reliability values on an individual subject basis (see Figure 3 and Supplementary Figure S1-S2).

**Figure 4.**
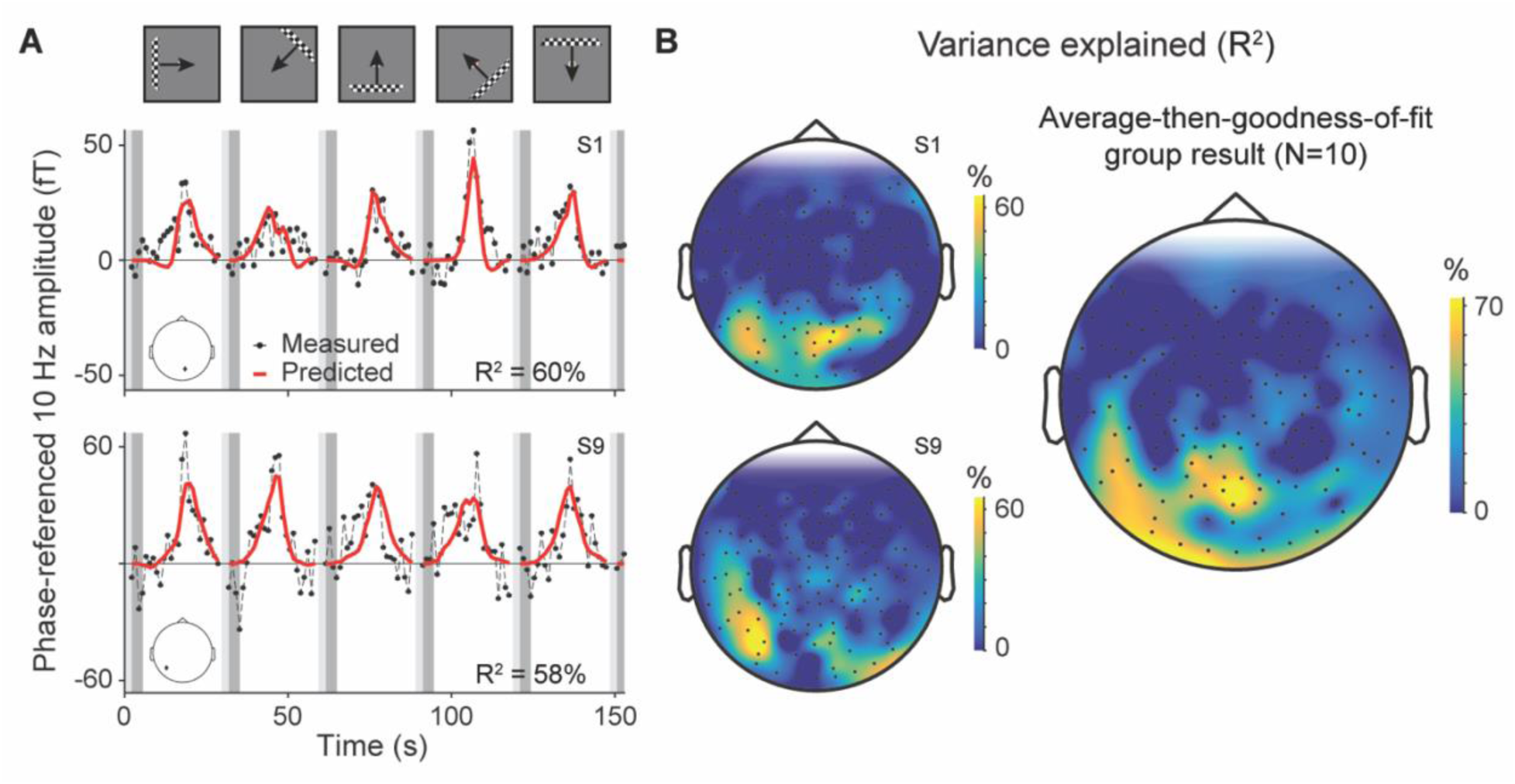
MEG forward model captures variance in observed MEG responses across posterior sensors. **(A)** Left panels show two example time series of observed 10 Hz phase-referenced MEG responses (black dots with dashed line) and predicted MEG responses by the model (red line). The predicted MEG responses explain 60% and 58% of the variance in the observed MEG responses. Data are from two posterior sensors (indicated by the black dot on the head schematic) in two different subjects (top: S1, bottom: S9). Every dot in the observed MEG time series is the phase-referenced 10 Hz amplitude of a single stimulus bar position. Light and dark gray boxes indicate blink and blank periods respectively. Blink periods were excluded from the analysis, blank periods were modeled as zeros. **(B)** Topographic sensor maps of variance explained by forward model. Left side shows the same two subjects as in panel (A) (top: S1, bottom: S9). Right side shows average-then-goodness-of-fit group result (N=10). In this case, measured MEG data are averaged across subjects and compared to the average across subject’s model fits.

One advantage of our forward model is that individual subject’s predicted MEG responses can be averaged to compare against the average observed data. We find that the average-then-goodness-of-fit group result could explain up to ∼70% of the variance in the average time series of several posterior sensors (Figure 4B, right panel). Because averaging across subjects’ data reduces measurement noise, the model fit is able to capture more variance in those sensors with high signal (posterior MEG sensors), compared to individual subjects. We also observe that the average-then-goodness-of-fit group result shows an asymmetry in captured variance explained, with higher variance explained on the left compared to right. However, it seems unlikely to reflect a general bias in the population, as individual subject maps do not support a systematic asymmetry in model accuracy between left and right sensors (Supplementary Figure S2). Rather, this more likely arises from better spatial alignment of sensors with good data on one side of the helmet than the other.

### 4.3 Forward model predictions are sensitive to changes in pRF parameters

Because MEG sensors pool over large regions of the cortex, the measured steady-state responses are the sum of many cortical pRF responses sampling visual space. This large pooling function poses the question: To what extent do the parameters of cortical pRFs in our forward modeling approach affect the accuracy of the predicted MEG responses? In the most extreme scenario, a forward model that uses scrambled pRFs across the cortex might predict MEG responses as well as the initially estimated pRFs. This would occur if each sensor pooled signals about equally from all of visual cortex. In this case, the MEG responses only contain information about stimulus onset and offset, not the specific spatial positions. A more likely possibility is that MEG sensor responses carry some information about the visual field position of stimuli, but at a lower spatial resolution compared to pRFs estimated by fMRI. In this case, it is an empirical question how much MEG sensor responses are affected by slight changes in underlying pRF models.

To quantify the extent to which our model accuracy depends on the measured pRF parameters, we artificially changed the pRF model parameters estimated from fMRI. First, we systematically alter pRF positions on the cortex, such that pRFs rotate around the fovea, leaving pRF sizes intact. Then, we systematically scale pRF sizes, leaving pRF positions intact. In both cases, we observe that the forward model predictions generally become less accurate.

#### 4.3.1 MEG data are best predicted by pRF positions estimated from fMRI

When rotating pRFs away from their estimated positions, the variance explained by the forward model decreases. For example, in subject S1 variance explained by the model decreased by ∼23% when rotating the pRFs from 90° clockwise or counter-clockwise around the fovea and slightly recovers when rotating 180° (Figure 5A, top panel). In other subjects, such as S9, variance explained peaked at the estimated pRF position, but the fall off with rotation angle was less steep (Figure 5A, bottom panel). For 3 out of 10 subjects (S1, S5, S6), variance explained by the model had a clear peak at 0° (the initial pRF position) and for 3 out 10 subjects (S4, S7, S9) variance explained peaked near 0° (±45°) (Supplementary Figure S3A). For the other 4 subjects, there was either a peak at an unexpected rotation (S3, S8), an asymmetric shape (S2) or a very small effect of pRF rotation (S10). On average, we observed the highest variance explained with 0° rotation, with a maximum drop of ∼15% when pRF positions were rotated around the fovea (Figure 5B).

**Figure 5.**
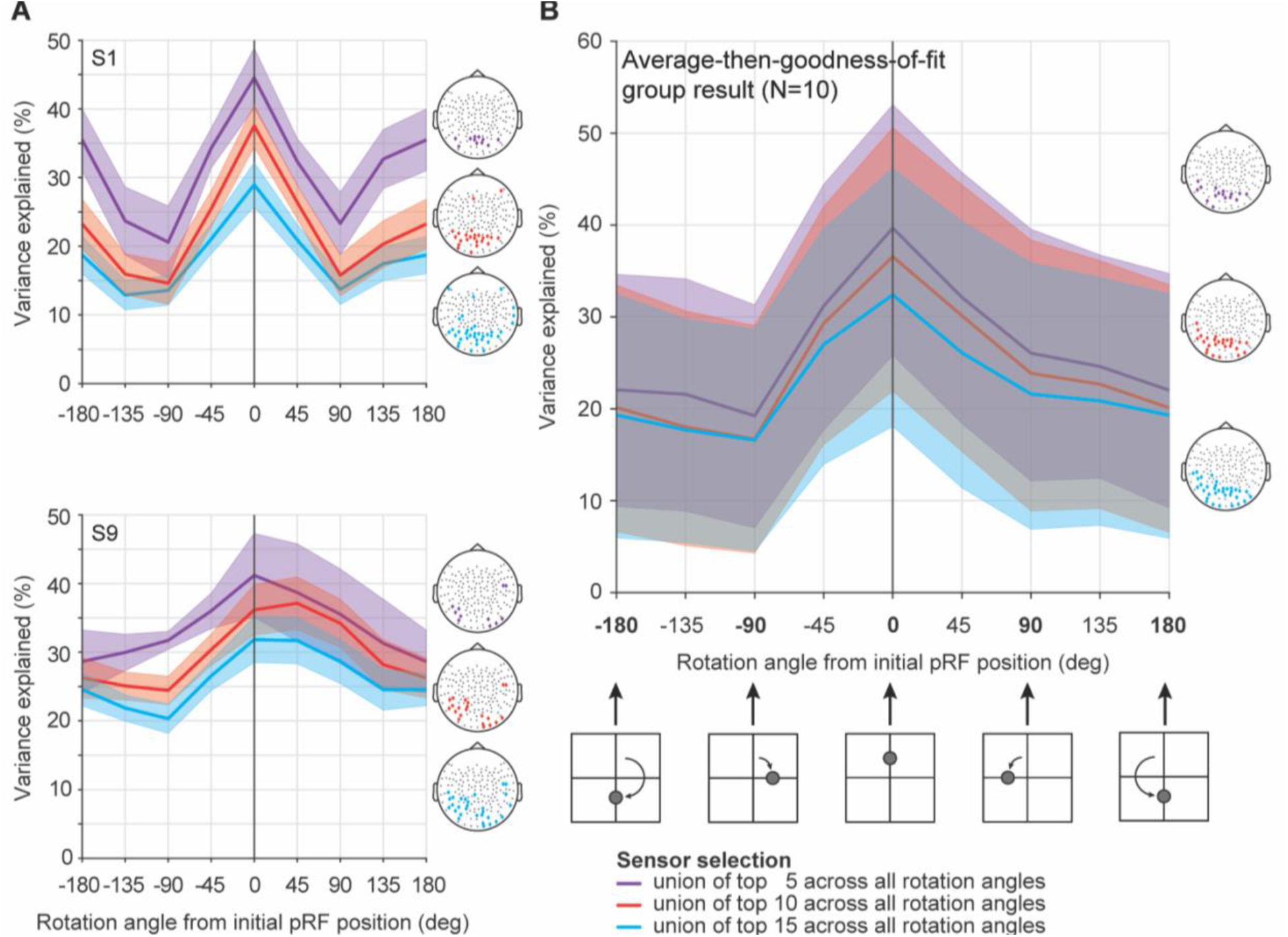
Systematic variation of pRF position decreases ability to explain variance in data by model predictions. **(A)** Variance explained by the forward model as a function of pRF center position for two subjects (top: S1, bottom: S9). The initial pRFs estimated using fMRI (0°, black vertical line) were systematically rotated around the fovea, by -180° to 180° from initial pRF position in steps of 45°. Predicted MEG responses were recomputed and fitted to observed MEG responses for each rotation condition. Data were summarized as the average across the union of 5 (purple line), 10 (red line), or 15 sensors (blue line) with the highest variance explained for each rotation condition (*i.e.,* including all sensors that are among the 5, 10, or 15 sensors with highest variance explained for at least 1 rotation direction; selected sensors are shown in schematic head plots on the right using the same color code). Shaded regions show ±1 standard error of the mean across the selected sensors. Highest variance explained is observed for the initial pRF position (0° rotation) for S1 for all 3 sensor selections and for S9 at the initial pRF position for top 5 sensors and near the initial position (between 0-45°) for top 10 and top 15 sensors. **(B)** Variance explained by average-then-goodness-of-fit group result and 68%-confidence interval (shaded region) obtained by bootstrapping 10,000 times the group average across 10 subjects for the sensor selection shown in schematic head plots on the left. Same color code is used as in panel A. A schematic of different rotation angles for an example pRF is shown below. On average, variance explained by the model fit decreases ∼15% when using pRF positions rotated away from the initial pRF position.

Rotating pRFs away from their estimated positions also affected the spatial topography of the predicted responses. When pRF positions were rotated away from their initial position, the sensors with the highest variance explained were confined to a single posterior region. The change in topography indicates that sensors differ in their sensitivity to pRF position (Supplementary Figure S4).

Importantly, the shape of the variance explained curve as a function of rotation angle does not depend on the exact number of sensors selected, although the overall variance explained decreases with the number of selected sensors. Averaging from only the top 5 sensors (purple line in Figure 5) results in the largest variance explained, and averaging from the top 15 sensors results in the lowest variance explained (blue line). This is expected because the more sensors that are included, the lower the average variance explained will be. The similarity in pattern as well as the difference in the mean as a function of the number of sensors included is found for both individual subjects (Figure 5A and Supplementary Figure S3A) and the average-then-goodness-of-fit group summary (Figure 5B). These analyses indicate that model sensitivity to pRF rotation is robust.

#### 4.3.2 Artificially changing pRF sizes affects model accuracy

When artificially altering pRF sizes 5x smaller or 10x larger, variance explained by the model gradually decreases up to 5-15%. We observed that our forward model explained on average most variance when using sizes close to, but slightly larger than, the pRF size estimated with fMRI (Figure 6). Some subjects showed a peak at slightly larger sizes (subject S1; Figure 6A, top panel), whereas other subjects had a local peak at slightly smaller pRF sizes (subject S9; Figure 6A, bottom panel). Overall, for 6 out of 10 subjects (S1, S3, S4, S5, S7, and S9) we observed a local peak in variance explained by the model at or near the initially estimated pRF (see Supplementary Figure S3B), most of them overlapping with those subjects showing a reliable effect of pRF position manipulation (see Supplementary Figure S3A). The other 4 subjects showed either a very small effect of scaling (S6), or the unexpected result of no effect for scale factors up to 1x and a monotonic increase in variance explained for scale factors larger than 1x (S2, S8, S10). We did not analyze scale factors beyond 10x to see if variance explained peaked for even larger pRF sizes, as those scale factors would make many pRFs extend beyond the stimulus field-of-view. When a pRF size becomes very large relative to the stimulus field-of-view, our experiment basically becomes an “on-off” paradigm with a full-field stimulus. These very large pRFs will therefore still capture some variance in our experiment. Indeed, with fMRI pRF models, a simple “on-off” model (*i.e.*, a non-spatially selective model which predicts a uniform response to a stimulus anywhere in the visual field relative to a blank) explains substantial variance in brain regions with very large pRFs (supplementary figure 11A in (Benson et al., 2018)). Hence, we expect that with scale factors beyond 10x the variance explained will decrease slightly but eventually plateau (and not go back to zero percent variance explained). See section 5.3 for further discussion of this observation.

**Figure 6.**
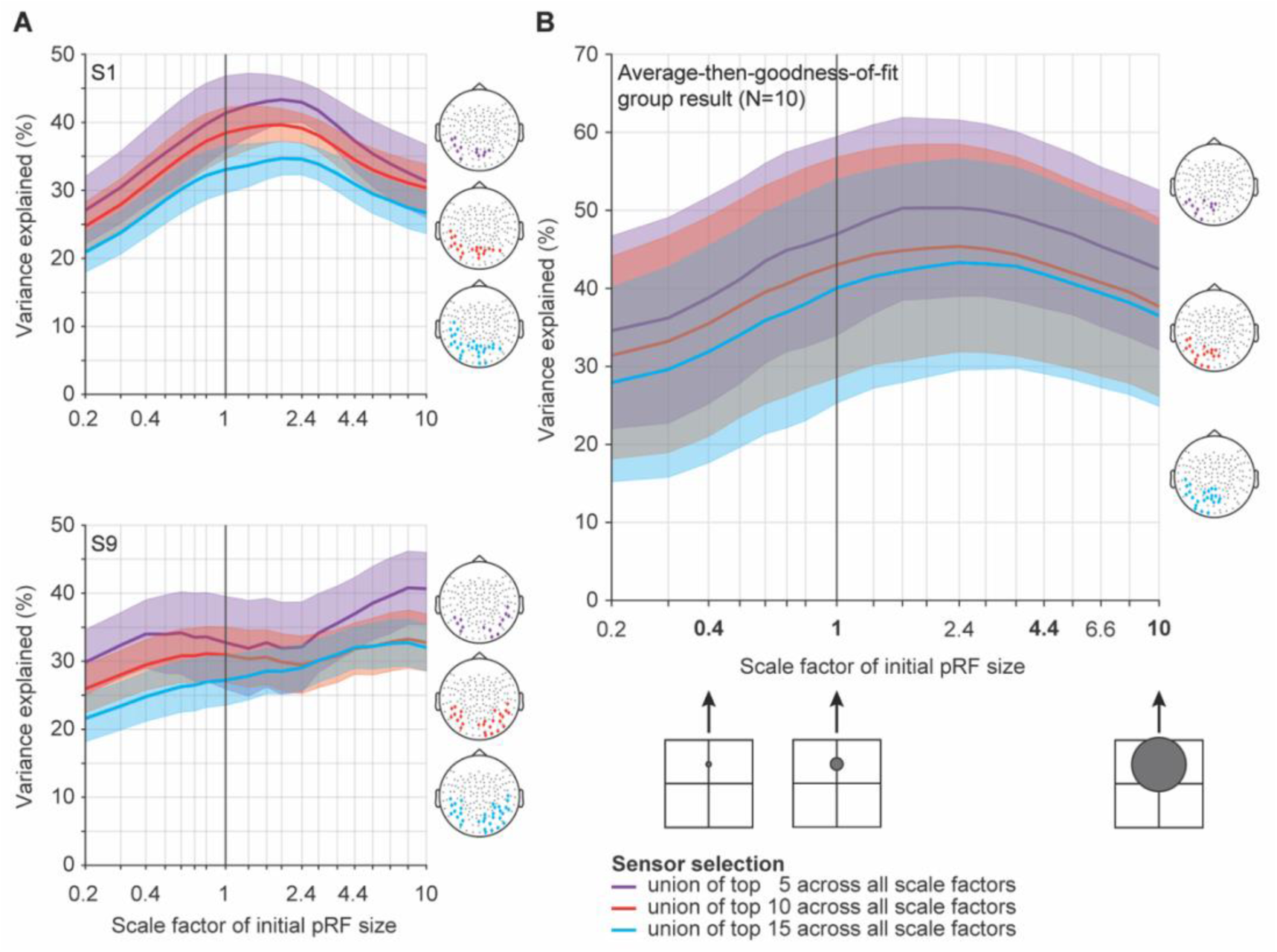
Systematic variation of pRF size decreases model accuracy. **(A)** Variance explained by the forward model as a function of scaled pRF sizes, *i.e.,* larger or smaller than initial pRF size estimated with fMRI (black line at 1). Top and bottom panels represent subjects S1 and S9, respectively. PRF sizes are systematically scaled from 0.2x to 10x the initially estimated size. Similar to variations in pRF position, variance explained is averaged across the union of 5 (purple line), 10 (red line), or 15 sensors (blue line) with the highest variance explained from each of the 19 scaling conditions. Shaded regions show ±1 standard error of the mean across the selected sensors. For S1, variance explained peaks at a pRF size that is close to, but slightly larger than initially estimated with fMRI for all 3 sensor selections. For S9, there is a local peak at a smaller size than initially estimated with fMRI using top 5 and 10 sensors, and then variance explained continues to increase at larger sizes. **(B)** Variance explained by average-then-goodness-of-fit group result for top 5, 10, and 15 sensor selection and 68%-confidence interval obtained by bootstrapping 10,000 times the group average across 10 subjects (shaded areas). Three schematic heads on the right show selected sensors for top 5, 10, 15 sensors using the same color scheme. Different scale factors for an example pRF are shown below the x-axis. On average, variance explained by the model fit decreases ∼15% when using pRF sizes that are 5x smaller or 10x larger than the initial pRF position.

Across subjects, we observed a similar drop in variance explained as a function pRF scale factor (around 15%), with a plateau between the initial pRF size and doubling the pRF size (Figure 6B). This indicates that MEG responses are less sensitive to changes in pRF size compared to pRF positions, or similarly, our forward model’s ability to capture pRF size changes. Changing the pRF size caused subtle changes in the spatial topography of the variance explained sensor map, but these changes were neither systematic nor large (Supplementary Figure S5).

The precise number of sensors used to summarize the model accuracy (5, 10, or 15) has little effect on the shape of the variance explained curves as a function of pRF scale factor (Figure 6, purple vs red vs blue line). However, as with the rotation analyses, the overall variance explained values decrease when adding more sensors to the selection. We find these patterns both for individual subjects (Supplementary Figure S3A) and for the group average (Figure 6B). This result is consistent with the pRF position variation results (Figure 5), showing that the results are robust to the exact number of sensors selected.

#### 4.3.3 Generalizability across methods of computing group average and selecting sensors

The group average results shown in Figures 5 and 6 reflect model accuracy for the sensor-wise averaged data. However, the sensors most responsive to the stimuli may differ across subjects. Therefore, we also computed group-average model accuracy by first summarizing the response for each subject as a function of pRF rotation or scale, and then averaging these response functions across subjects (goodness-of-fit-then-average). This method uses the best sensors for each subject, either as defined previously (highest variance explained) or defined in a model-independent manner (highest split half reliability of the 10 Hz SSVEF), and therefore respects individual differences in sensor topography. For individual subject data using the split-half reliability method for sensor selection, see Supplementary Figure S6.

The results from these analyses are similar to those we observed previously from the average-and-then-fit results. For pRF position variations, the variance explained curves peak at 0°, declining systematically when rotating away from the initial estimated pRF position (Figure 7A). For systematic variations in pRF size, the variance explained curves show a rise from reduced pRF sizes (0.2x) to the initially estimated pRF size (1x) (Figure 7B), similar to the average-then-goodness-of-fit results. The curves differ slightly from the goodness-of-fit-then-average results at higher scaling values, either plateauing or very slightly declining. These results further support the finding that the model accuracy is sensitive to pRF size and position parameters.

**Figure 7.**
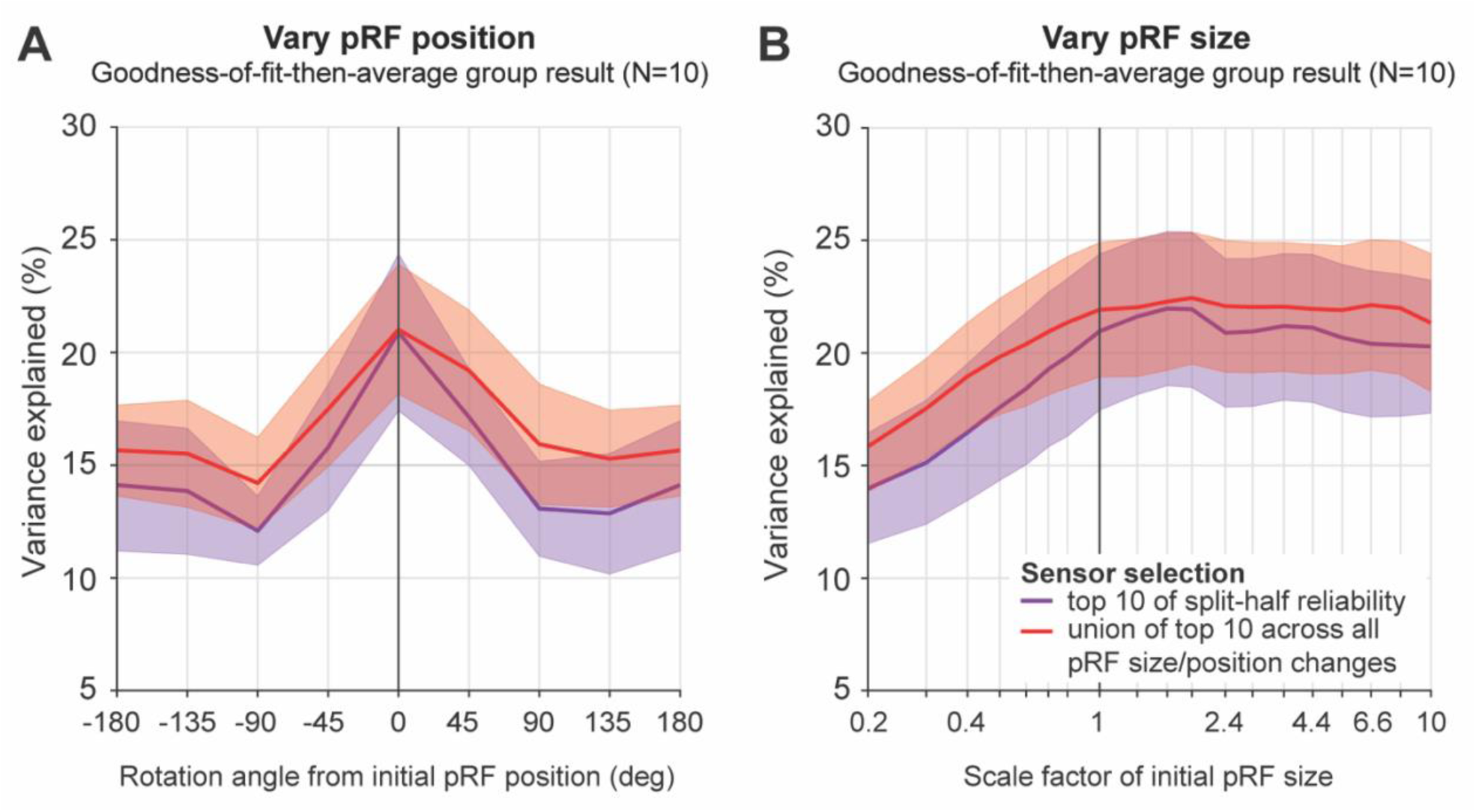
Group average effect of fitting individual data first, before averaging across subjects, using a model-based and data-based sensor selection. **(A) Effect of systematically varying pRF position.** Group average is computed by taking each subject’s variance explained curve and averaging across subjects at each rotation angle or scale factor. Data were bootstrapped across subjects (10,000 times), where lines represent the average across bootstraps and shaded areas 68%-confidence intervals. Red colors represent results using a model-based sensor selection (for each subject, the union of top 10 sensors across all pRF position variations). Purple colors show results using a data-based sensor selection (for each subject, the 10 sensors with the highest SSVEF split-half reliability). Individual subject data and schematic head plots are shown in Supplementary Figure S6. **(B) Effect of systematically varying pRF size.** Same color code as in panel A, but now for bootstrapped goodness-of-fit-then-average group result when systematically scaling pRF sizes.

While the shape of these functions is similar to those in Figures 5 and **6**, the overall height (*i.e.*, mean variance explained values) is lower. This is because the average-then- goodness-of-fit method reduces uncorrelated noise in the measurements, allowing for a higher variance explained by our model if its underlying assumptions are correct. In contrast, the goodness-of-fit-then-average method by definition preserves the average variance explained.

## 5 Discussion

Population receptive field modeling is an important tool that has made significant contributions to our understanding of the functional architecture and underlying computations of the human visual cortex. The successes of pRF models have been widespread and large in fMRI, with a few applications in intracranial data, and little applications for MEG forward models. Here, we developed a stimulus-to-sensor forward model that combines pRFs estimated from fMRI with a biophysical forward model to predict the steady-state visually evoked MEG responses when subjects viewed moving bar stimuli. Our results demonstrate that we can reliably measure and predict visually-evoked responses for these stimuli. The model was sensitive to cortical pRF model parameters, as we found a decrease in variance explained when artificially changing the underlying pRF model parameters estimated with fMRI.

This combination of fMRI and MEG measurements allows future studies to investigate the time-resolved spatiotemporal dynamics of human visual field maps as well as the relationship between the fMRI BOLD response and electromagnetic field measurements. In principle, our forward model can be implemented without solving pRF models using fMRI data. This can be done by applying a retinotopic template to an anatomical MR image, for example (Benson et al., 2012; Benson, Butt, Brainard, & Aguirre, 2014; Benson & Winawer, 2018), or by predicting retinotopic structure from the cortical curvature pattern via machine learning algorithms (*e.g.,* deep neural networks (Agrawal, Stansbury, Malik, & Gallant, 2014; Khaligh-Razavi & Kriegeskorte, 2014; Güçlü & van Gerven, 2015; Eickenberg, Gramfort, Varoquaux, & Thirion, 2017; Güçlü & van Gerven, 2017; Ribeiro, Bollmann, & Puckett, 2020)). Such applications would simplify and shorten the solution to the model parameters and reduce MRI scanning time which is useful when studying special populations like children or patients or individuals having difficulty holding fixation.

As a proof of principle, we implemented a forward model that uses group average pRF parameter maps from an aggregate retinotopy dataset (Himmelberg, Kurzawski, et al. (2021)) and compared its model performance to our standard forward model using subject specific retinotopy data. These average pRF maps were collected with stimuli of approximately the same field-of-view but differing in pattern. The overall model accuracy was similar to that obtained from using pRFs measured in individual subjects (Supplementary Figure S7A). Moreover, the specific variance explained topographic sensor maps were broadly similar for the two methods (a correlation of about 0.6 between sensor maps from the same participant across the two methods, compared to about 0.2 for different participants; Supplementary Figure S7B). This result shows that average pRF parameter maps from an aggregate retinotopy dataset can be used to make reasonable forward model predictions.

In contrast, it may be more problematic to use an average anatomical template and head model to build cortical predictions with our forward model, rather than using the subject’s measured head anatomy and head position. This is because the cortical folding geometry of an average anatomical template is not realistic; it emphasizes large sulci and gyri and removes idiosyncratic folding patterns of individual subjects. Moreover there are large differences in the shape and size of visual areas, differing by as much as 3:1 across people (Dougherty et al., 2003; Benson et al., 2021). These differences are likely why we found higher pairwise correlations comparing variance explained sensor maps of the two forward models within subjects compared to across subjects (Supplementary Figure S7B).

### 5.1 Relationship to reconstructing cortical retinotopy from MEG sensor responses

Several MEG studies have aimed at reconstructing retinotopy responses on the cortical surface from MEG sensor measurements (*e.g.* (Moradi et al., 2003; Poghosyan & Ioannides, 2007; Sharon, Hämäläinen, Tootell, Halgren, & Belliveau, 2007; Brookes et al., 2010; Perry et al., 2011; Cicmil, Bridge, Parker, Woolrich, & Krug, 2014; Nasiotis, Clavagnier, Baillet, & Pack, 2017)). In those studies, instead of a forward model from stimulus to sensors, the cortical sources are estimated by inverse modeling: going from sensors to cortical sources, that is, estimated sources are derived by multiplying the sensor responses by the pseudo-inverse of the gain matrix in the head model. These estimated source responses are then compared to visual field maps measured with fMRI, where the fMRI maps are assumed to be the ‘ground truth’, aiming to minimize localization error.

This inverse modeling approach can localize the retinotopic responses within a centimeter on the cortex of the correct hemifield, but it is limited to early visual areas and fails to accurately capture known features of visual field maps. For example, stimuli in the upper visual field (*i.e.,* the lower bank of the calcarine sulcus) cannot be captured due to low SNR or signal cancellation in MEG sensors (*e.g.,* see (Nasiotis et al., 2017)). Additionally, changes in stimulus polar angle and eccentricity—a hallmark of visual field maps—can only be distinguished at a coarse scale (*i.e.* visual quadrants or fovea versus periphery) (Moradi et al., 2003; Brookes et al., 2010; Perry et al., 2011; Cicmil et al., 2014). One reason for these limitations is that the inverse problem is ill-posed: a measured magnetic flux from a single sensor can result from an infinite number of cortical source combinations. The solution to this inverse problem is ill-defined and can only be achieved by making assumptions to limit the possible solutions (Cicmil et al., 2014). Several research groups have used the known location of visual field maps as a prior to constrain the number of possible solutions, also known as ‘Retinotopy Constrained Source Estimation’ (Hagler et al., 2009; Ales, Carney, & Klein, 2010; Hagler & Dale, 2013; Hagler, 2014; Cottereau, Ales, & Norcia, 2015; Inverso et al., 2016). These constraints resolved some of these reconstruction errors (*e.g.,* cross-talk between sources in visual areas with close proximity, see (Hagler et al., 2009; Cottereau et al., 2011; Cottereau et al., 2015)), but the overall approach of source reconstruction still relies on regularizers coming with certain assumptions.

Our forward model takes a different approach from previous MEG studies: we turn inverse modeling on its head. With our approach, model predictions are not limited to early visual areas, but only by the extent of reliably estimating local pRFs on the cortex. Also, our approach is not constrained by cancellation effects of opposite facing dipoles. On the contrary, our approach can be used to investigate the effect of source cancellation on sensor responses by simulating different temporal patterns in visual cortex (Kupers, Benson, & Winawer, 2020). We first predict neural time series at a millimeter-scale on the cortical surface using local pRF models estimated with fMRI, before predicting sensor responses with the MEG forward model. Because we use a purely forward modeling approach, our model is well-defined and avoids the need for additional constraints. By introducing an intermediate step, *i.e*., modeling responses on the cortical surface, between the stimulus and the MEG sensor responses, our model has the ability to implement a quantitative description of the stimulus representation at the cortical source level; information one usually does not have access to and aims to reconstruct. Because our model is informed by local pRFs, it can create predictions at the millimeter scale, hence incorporating stimulus-selectivity at a local scale, and thereby make meaningful and accurate predictions at the MEG sensor level.

In addition, having a computational encoding model that predicts sensor responses at an individual subject level introduces an alternative way of summarizing group data. Instead of computing sensor-wise average of the summary statistic (for example, variance explained), it is possible to average individual data and individual predictions separately and compare the average group prediction to the average group data (“average-then-goodness-of-fit”). Typically, MEG or EEG sensor data averaged across subjects can be difficult to interpret. Because of individual differences in cortical geometry and head position, a particular sensor will pool over different brain sources from each subject. For this reason, the sensor-wise averaged data are not easily linked in a meaningful way to cortical sources or to the experimental paradigm. In our case, however, the forward model for each subject respects the variation in pRF parameters across that subject’s cortex, as well the subject’s cortical folding pattern and head position in the MEG helmet. Hence the averaged model predictions, though summarized in the sensor space, reflect details of each of the individual subjects, and provides a compact summary of the result. Unlike averaging over, say, repeated trials within an individual, averaging over subjects entails some degree of uncorrelated signal (due to the differences in subject cortical geometry) in addition to uncorrelated noise. Hence the SNR is not expected to increase in a simple way as the number of subjects increases. Nonetheless, the SNR is higher for the average-then-goodness-of-fit method than for any individual subject.

Nonetheless, the average-then-goodness-of-fit method summary has some interpretation limits. For example, it may result in a smoother topographic map than is found for any individual and will tend to show more accurate predictions in locations where the topographic maps are better aligned across subjects. For these reasons, we confirmed our results with the goodness-of-fit-then-average method, which shows lower variance explained, but respects differences between subjects in terms of which sensors show the best model fits.

### 5.2 The relationship between MEG and fMRI measurements

MEG and fMRI are two of the most widely used non-invasive measurement techniques in human neuroscience capturing different types of aggregated responses across neural populations. MEG captures the magnetic flux from local field potentials, whereas fMRI captures the neurovascular response. The neural signals giving rise to each measurement are likely to differ. For example, the MEG signal is most sensitive to pyramidal neurons whose dendrites are perpendicular to the cortical surface (Hämäläinen, Hari, Ilmoniemi, Knuutila, & Lounasmaa, 1993), which may differ from sensitivity of the fMRI BOLD signal. Moreover, the neural signals giving rise to the fMRI signal have been shown to be most similar to those giving rise to the broadband component of the field potential, not the evoked signal which we used here (Foucher, Otzenberger, & Gounot, 2003; Winawer et al., 2013; Hermes, Nguyen, & Winawer, 2017). These factors will put an upper limit on how well our model can perform. Nonetheless, differences in tuning of the neural populations giving rise to different signals are likely to be modest in the domain of position tuning, considering that position tuning is mapped at a relatively large scale in cortex (millimeter), compared to other features such as orientation, eye of origin preference, or spatial frequency preference, which may vary at a finer spatial scale.

### 5.3 Sensitivity differences in predicting pRF position and size for fMRI vs MEG

We showed that when artificially rotating pRF positions on the cortical surface, the model explained most variance in the data for the pRF positions obtained by fMRI. This indicates that the optimal pRF position explaining fMRI BOLD data also predicts the steady-state responses best in MEG sensors. On the other hand, artificially scaling pRF sizes did not cause our model performance to peak at the estimated pRF size. For several subjects, we observed a local peak in variance explained for models using pRF sizes slightly larger, while others for slightly smaller, than those estimated from fMRI.

Given that we observed 10 Hz steady-state amplitudes with high reliability and signal-to-noise ratio in posterior MEG sensors, it is unlikely that the differences between data and model predictions are solely caused by measurement noise. In addition, our model is fairly conservative and unlikely to overfit MEG data as it contains relatively few free parameters (one gain factor and one reference phase per MEG sensor) which undergo a cross-validation procedure.

In terms of modeling, the pRF size discrepancy can arise if the initial fMRI estimates overpredict pRF size, our MEG forward model underpredicts pRF size, vice versa, or a combination of both. Several neural and non-neural factors have been reported to bias estimated pRF sizes with fMRI, whereas pRF position estimates appear to be more robust.

### Non-neural factors

One non-neural factor that has a large effect on the estimated pRF size (and less so for pRF position) is the mismatch between the assumed and actual underlying hemodynamic response function (HRF). This mismatch can cause both over- and underestimation of pRF sizes, depending on the experimental design or whether the spatial or temporal component of the assumed HRF is inaccurate (Dumoulin & Wandell, 2008; Lerma-Usabiaga, Benson, Winawer, & Wandell, 2020). Since our fMRI session used stimuli that swept across the visual field in both directions for a given orientation, we believe that our experimental design minimized any bias in the estimated pRF size caused by the sluggish HRF. We did not estimate HRF functions separately for individual subjects or visual areas. We also did not model the spatial component of the HRF. However, our presentation time of sweeping bars was relatively long (31s/bar sweep), which largely reduces the impact of pRF size biases caused by the HRF mismatch (Lerma-Usabiaga et al., 2020).

Another possible non-neural factor that has been reported to bias pRF sizes are eye movements. As shown by simulation (Levin, Dumoulin, Winawer, Dougherty, & Wandell, 2010; Klein, Harvey, & Dumoulin, 2014) and empirically (Hummer et al., 2016), gaze instability can introduce overestimation of pRF sizes across eccentricity. It also increases the absolute mean error for pRF position, but with no systematic bias within polar angle or eccentricity maps compared to gaze-corrected fMRI data. In the present study, eye movements were monitored during fMRI and MEG experiments for most subjects and did not show large eye movements. However, we cannot rule out the presence of small fixational eye movements (*i.e.,* microsaccades and drift) in both MRI and MEG sessions. At least, if microsaccades were present in the MEG data they would not cause an electromagnetic field response that overlaps with the 10 Hz steady-state response, as microsaccades are reported as increased gamma-band power (> 60 Hz) (Yuval-Greenberg, Tomer, Keren, Nelken, & Deouell, 2008).

#### Neural factors

A neural factor that could affect pRF properties is visuo-spatial attention. FMRI and MEG sessions contained the same stimuli and similar experimental design where subjects were performing a fixation task. However, we cannot rule out fluctuations in covert spatial attention shifts towards the moving bar stimulus (either voluntary or involuntary). Several fMRI studies that explicitly manipulated voluntary visuo-spatial attention reported changes for pRF positions, and no changes or much less so for pRF sizes (Klein et al., 2014; Kay, Weiner, & Grill-Spector, 2015; Vo, Sprague, & Serences, 2017; van Es, Theeuwes, & Knapen, 2018). While individual subjects could employ different amounts of visuo-spatial attention in one session compared to the other, on average our initial estimates of pRF position seem more robust compared to pRF size. This suggests that visuo-spatial attention is unlikely the main factor causing a difference in optimal pRF size for MEG versus fMRI.

### 5.4 Choice of MEG data component

In this study, we compared the phase-referenced steady-state amplitudes against the predicted retinotopy response. We chose SSVEFs because this signal contains stimulus-specific information (*i.e.,* the contrast-reversal rate) and has a high signal-to-noise ratio. However, we do not exclude the possibility that other MEG data components are a better proxy for the predicted retinotopy responses in MEG sensors.

Our model predictions are based on local pRFs estimated from fMRI BOLD responses, but the measured SSVEFs originate from high coherence between neural sources—a signal type fMRI is less sensitive compared to electric field measurements like ECoG (Foucher et al., 2003; Hermes et al., 2017). For example, the ECoG study by Winawer *et al*. (2013) used a similar experimental design as the current study: presenting high contrast-reversing checkerboard bars traversing across the visual field while recording local field potentials from early visual cortex. They found that when a bar crossed the estimated pRF of the ECoG electrode, there was an increase in steady-state amplitude at the stimulus frequency *and* a broad increase in power across many frequencies, *i.e.,* a parallel shift of the 1/f spectrum compared to baseline (“broadband response”). When comparing both data components to BOLD responses of pRFs at the same cortical location in healthy controls, the broadband response was a better predictor of spatial summation compared to the steady-state response. This difference becomes clear when using test stimuli that vary in bar width or size. In this case, both the fMRI and the broadband signal show sub-additive summation, whereas the evoked response does not. Had we used stimuli with multiple bar widths and sizes in our MEG experiment, model accuracy for the SSVEF would likely have been lower.

### 5.5 Choice of model parameters

Currently, our model predicts responses from stimulus to cortex without free parameters (after the pRF models are solved for fMRI) and fits two free parameters per MEG sensor (a reference phase and gain factor). Using a limited number of free parameters makes our model predictions interpretable: the reference phase allows for a sign reversal of the MEG prediction and potential delays in visual processing across the visual hierarchy, and the gain factor puts the model predictions in units of femto-Tesla. Allowing additional free parameters (such as an offset or scale factor for pRF estimates on the cortex) or refitting our gain factor to the average of all MEG data runs is likely to improve model performance but can also cause overfitting or reduce its interpretability.

Additionally, other encoding models predicting visual preferences of neural populations could capture more complex dynamics compared to the current model. Examples of such models are the difference of Gaussian (DoG) pRF model (Zuiderbaan, Harvey, & Dumoulin, 2012) or the compressive spatial summation (CSS) model (Kay, Winawer, et al., 2013). Since our model implements the step from stimulus to predicted cortical responses in a separate function, the model component can be interchangeable and allows the general modeling approach to adapt to different experiments.

### 5.6 Individual differences

We observed that the amount of variance explained by our model was considerably different across subjects, using both the originally estimated pRFs with fMRI and when artificially varying pRF size or position. This inter-subject variability could be the result of methodological errors, measurement noise, non-neural physiological noise (such as head and eye movements), or a true difference between subjects. Methodological errors include the possibility of improper alignment of the MEG sensor positions to subject anatomy, and the type and resolution of the head model.

MEG and EEG head models have become increasingly more complex (for an overview, see (Vorwerk et al., 2014)). For example, we used the overlapping spheres method (Huang et al., 1999), but there are more biologically accurate models like the boundary element method (‘BEM’, (Kybic et al., 2005; Gramfort et al., 2010)). With the head model we used, we explained up to about 60% of variance in the sensor data. This is relatively close to the about 80% split-half reliability of the 10 Hz steady-state response, a proxy for the noise ceiling. Nonetheless there remains unexplained variance, indicating that there is some room for higher accuracy from better methods. The current approach provides a clear proof of principle that a forward model from stimulus top sensors can accurately predict responses to visual stimuli.

### 5.7 Future applications and extensions

Our forward model shows that MEG responses can be reliably predicted from stimulus to cortex to sensors. One interesting potential application to use our model is to characterize the changes in pRF properties over time. As mentioned previously, several fMRI studies have observed changes pRF center of mass with visuospatial attention (Klein et al., 2014; Kay et al., 2015; Vo et al., 2017; van Es et al., 2018). Our MEG forward model could be used to predict these changes and capture the time-resolved effects of visuo-spatial attention. A second application of our model would be the combination of spatial pRF models estimated with models that capture pRF preferences in temporal processing (Stigliani, Jeska, & Grill-Spector, 2017; Zhou, Benson, Kay, & Winawer, 2018) or replace the local pRF models on the cortex with topological maps coding for other types of perception (such as audition (Saenz & Langers, 2014)), cognition (such as numerosity (Harvey, Klein, Petridou, & Dumoulin, 2013)) or action (Mattay & Weinberger, 1999).

Future studies can extend our forward modeling approach and apply it to study a variety of questions aiming at spatiotemporal dynamics of visual processing. For example, one consideration is changing the experimental design of the MEG session. In the current study, MEG stimuli were designed such that they were similar to the retinotopic stimuli used for fMRI studies. However, because fMRI experiments sample BOLD responses at second time resolution and need to take into account the sluggish hemodynamic response, it does not mean that MEG measurements need to be sampled at the same time resolution with the same temporally predictable stimulus sequence. Since our model predicts the MEG responses to arbitrary stimulus apertures in the visual field based on the cortical spatial tuning preferences, it can predict other temporal sequences and give insight to a variety of spatiotemporal dynamics at sub-second temporal resolution.

### 5.8 Conclusion

Neuroscientists use a number of techniques to measure neural activity, each providing different information about brain activity. MEG measures the magnetic field induced by electric currents present in neural activity, whereas fMRI measures the metabolic demands associated with neural activity. In this paper, we demonstrate a forward model that can capture MEG sensor responses to retinotopic mapping stimuli, by combining pRFs estimated from fMRI responses with the biophysical MEG head model. Our results support a common underlying mechanism of neural processing measured with the two modalities, and provide new opportunities to study time-resolved spatiotemporal dynamics in visual processing.

## Author contributions

**ERK**: Methodology, Formal analysis, Investigation, Writing - Original Draft, Writing - Review & Editing, Visualization. **AE**: Methodology, Formal analysis, Visualization, Writing - Original Draft. **NCB**: Methodology, Supervision. **WZ**: Supervision. **MCJ**: Supervision. **SOD**: Conceptualization, Funding acquisition, Supervision. **JW**: Conceptualization, Methodology, Funding acquisition, Supervision.

## Acknowledgements

This project has received funding from the NIH Brain Initiative R01 MH111417-0 (JW), European Union’s Horizon 2020 research and innovation programme under the Marie Sklodowska-Curie grant agreement No 641805 (SOD), Ammodo KNAW Award (SOD) and NWO-VICI grant 016.Vici.185.050 (SOD). We thank Barrie Klein for making contributions early in the project to experimental design, data collection, and analysis.

## Competing interests

The authors declare no competing interests exist.

## 7 Supplementary figures

**Supplementary Figure S1.**
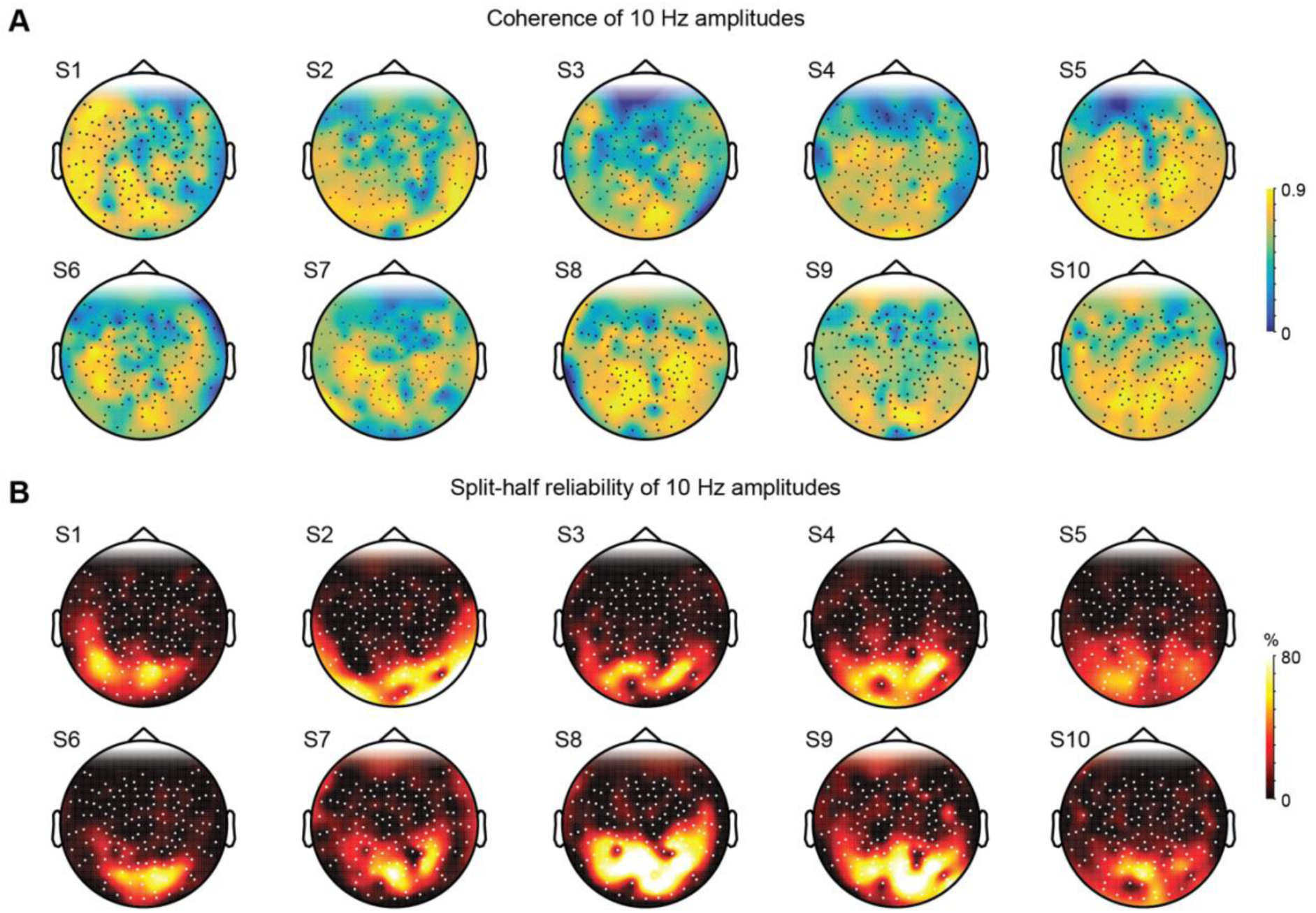
Coherence and split-half reliability of individual subjects of 10 Hz SSVEF amplitudes. **(A)** Coherence metric is computed as the amplitude of 10 Hz divided by the sum of 9 to 11 Hz. All subjects are plotted with the same color bar limits shown on the right. **(B)** Split-half reliability is calculated as the mean Pearson’s *ρ* correlation of 10 Hz amplitudes across 1000 iterations. Amplitudes within a single iteration are averaged across runs within each split-half and not phase-referenced. All subjects are plotted with the same color bar limits shown on the right.

**Supplementary Figure S2.**
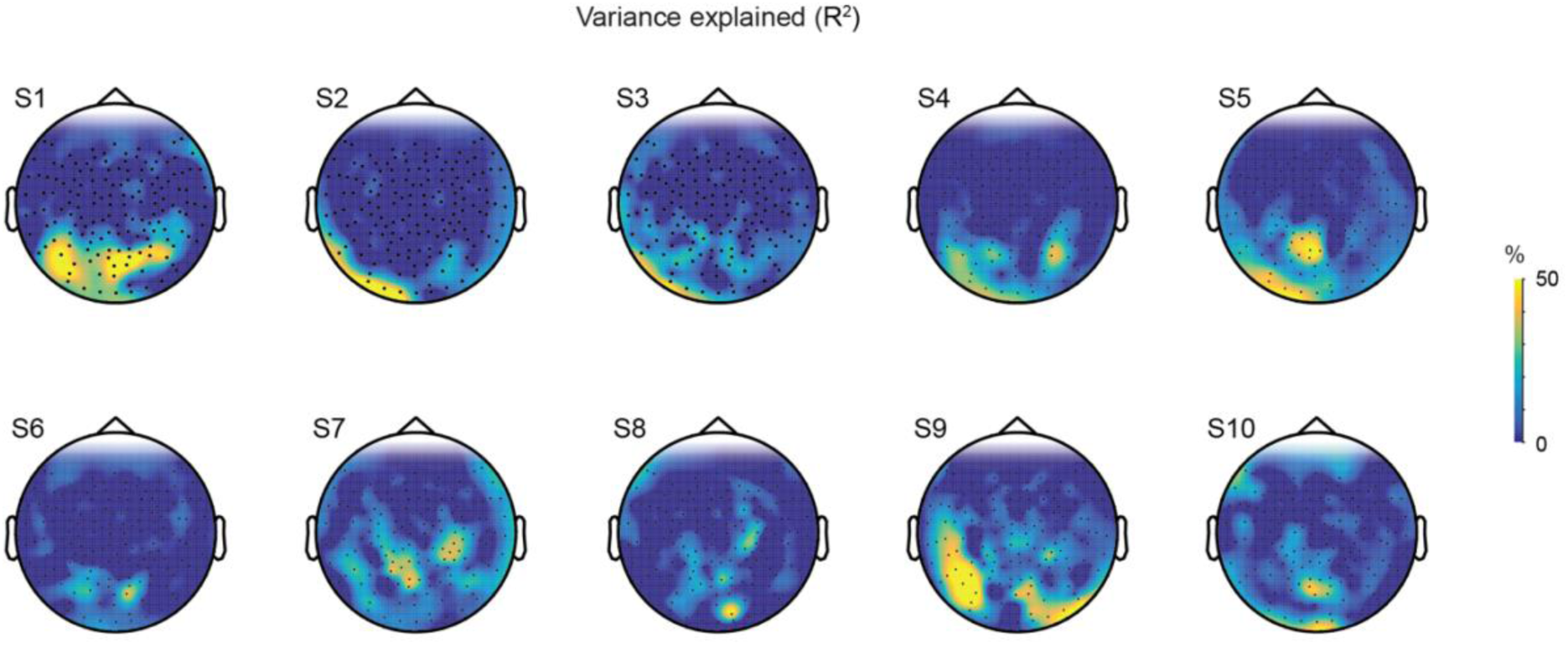
Topographic MEG sensor maps of variance explained by the model for all 10 individual subjects. The model is able to predict the measured 10 Hz phase-referenced steady-state MEG responses up to 50% of the variance in the measured MEG responses in posterior sensors for many subjects.

**Supplementary Figure S3.**
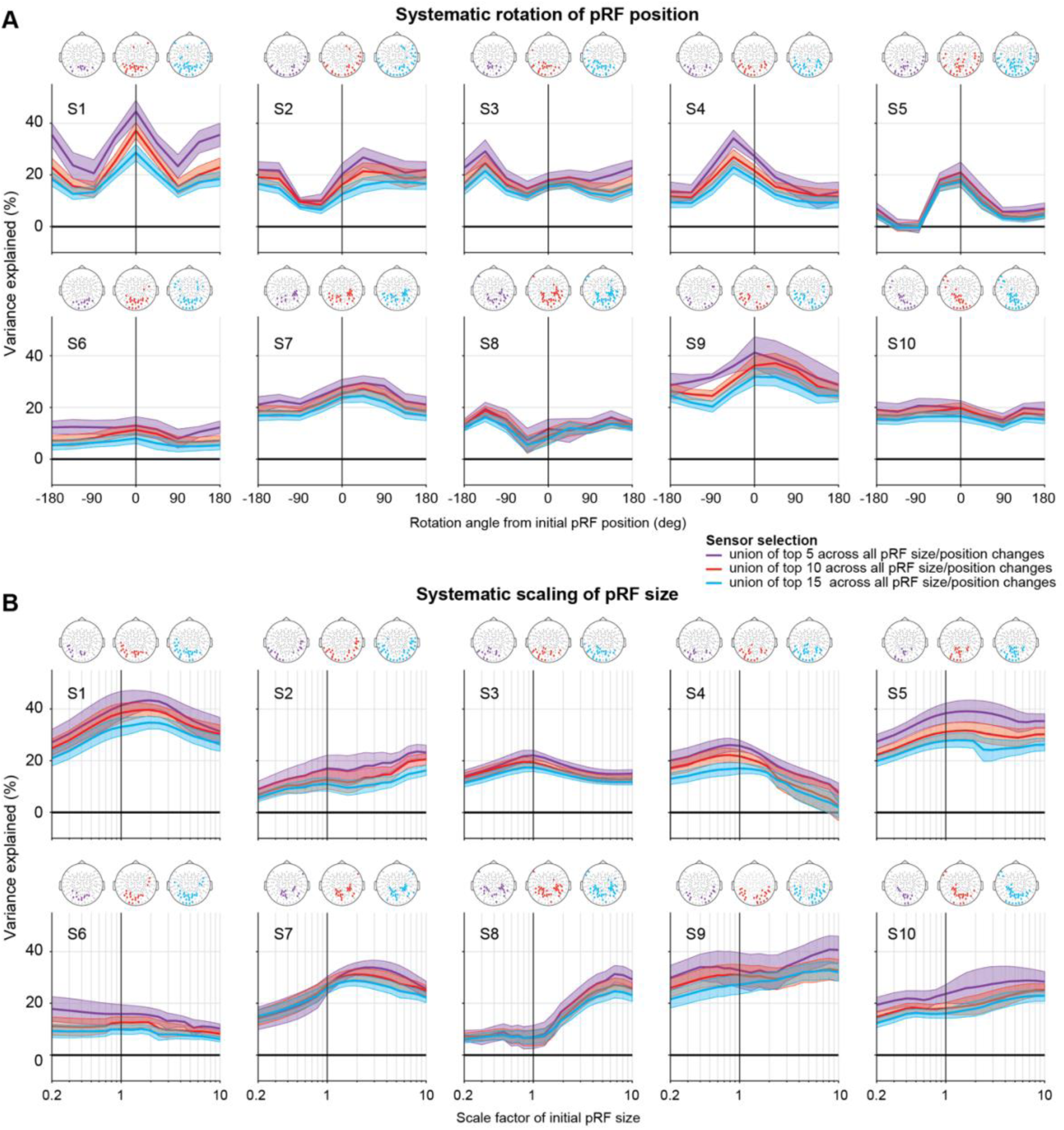
Variance explained by the forward model as a function of systematic pRF parameter changes for individual subjects. **(A) Effect on pRF position rotation.** The pRFs estimated with fMRI were systematically rotated around the fovea, by -180° to 180° from their original position in steps of 45°. Predicted MEG responses were then recomputed for each of the rotation conditions. Per subject, highest variance explained values were averaged from the union across the top 5 (purple), top 10 (red), or top 15 sensors (blue) in each of the 9 rotation conditions (dots in schematic head, same color scheme). Shaded regions show ±1 standard error of the mean across the selected sensors. While there are large individual variations, 6 out of 10 subjects (S1, S4, S5, S6, S7 and S9) have most variance explained in the MEG data when the initial pRF positions or near initial (±45 deg) were used in the forward model. **(B) Effect on pRF size scale factor.** Same as (A) but now for size scale factor variations. PRF sizes were systematically scaled from 5x smaller to 10x larger the initial size. Although there are large variations between individual subjects, 6 out of 10 subjects (S1, S3, S4, S5, S7, and S9) showing a local peak in variance explained for pRF sizes that are slightly smaller or larger than the original pRF size (a scaling factor of 1). In some, but not all subjects this local peak is followed by an increase in variance explained for very large-scale factors.

**Supplementary Figure S4.**
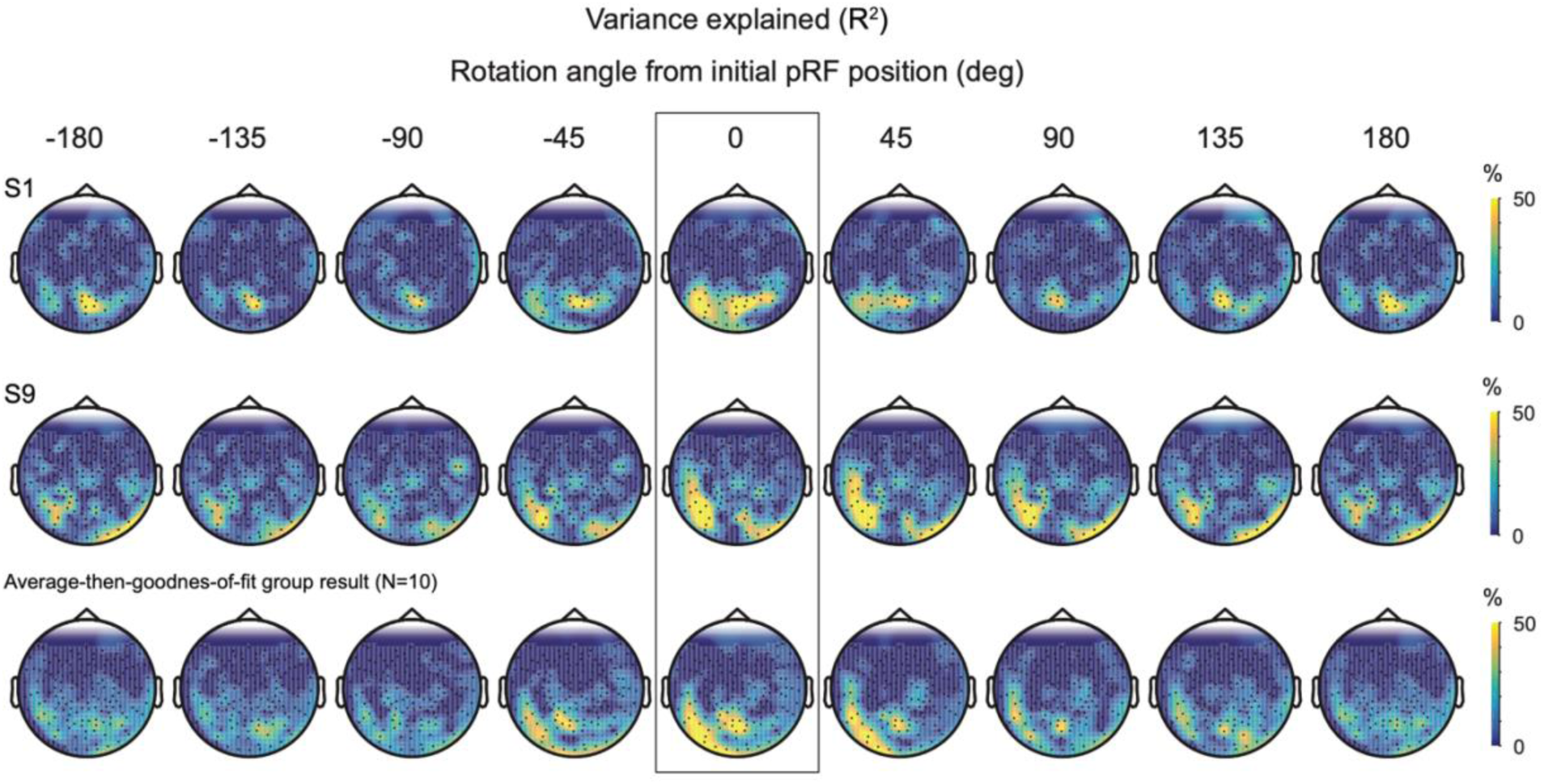
Topographic maps of variance explained by the forward model for different rotation angles of the initially estimated pRF position. Top and middle row show maps for 2 individual subjects (S1 and S9). Bottom row shows average-then-goodness-of-fit group result. Initial pRF position estimated by fMRI is outlined (0°). All maps use the same color scale as shown on the right.

**Supplementary Figure S5.**
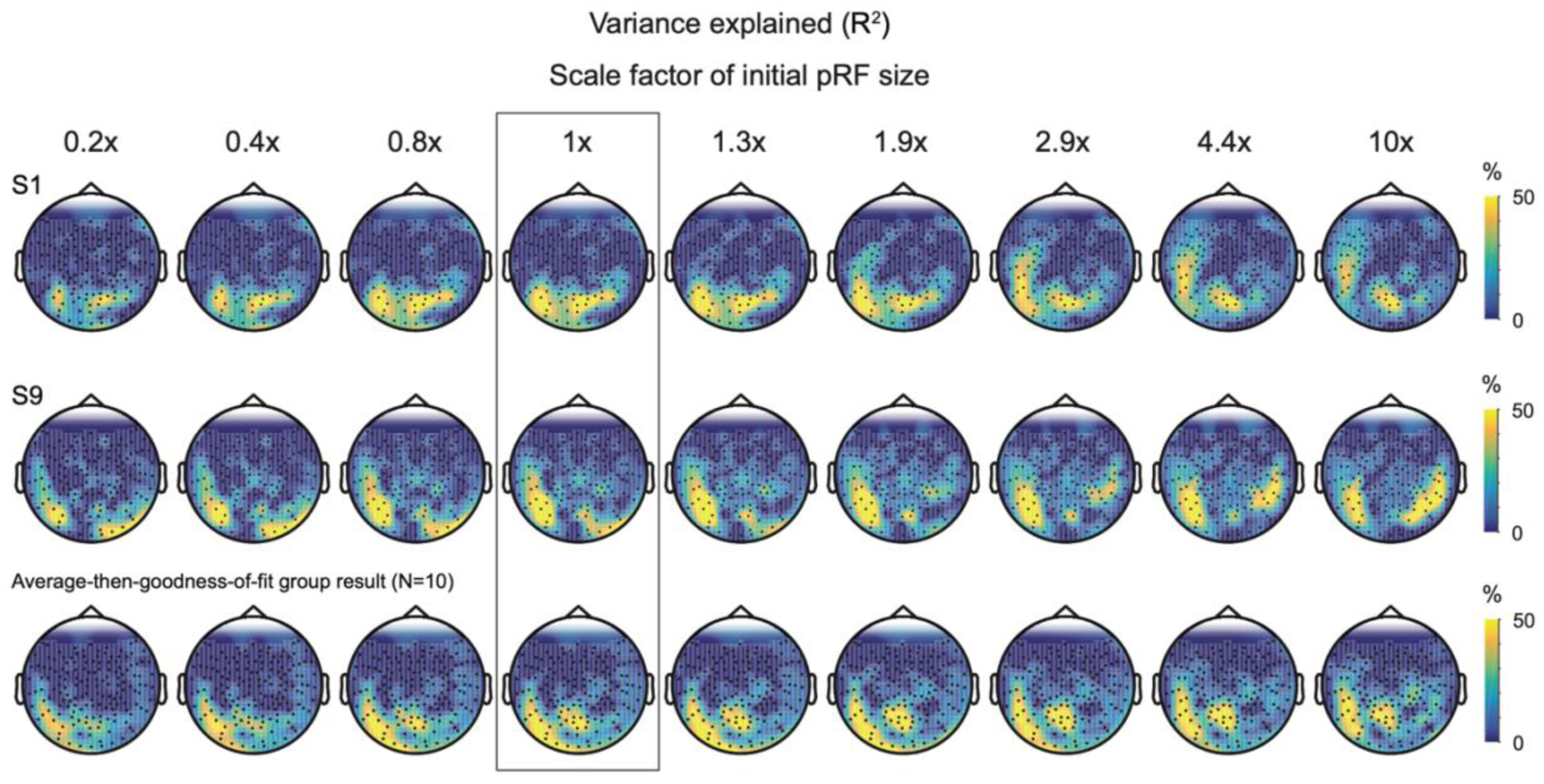
Topographic maps of variance explained by the forward model for different size scale factors of the initially estimated pRF size. Top and middle row show maps for 2 individual subjects (S1 and S9). Bottom row shows average-then-goodness-of-fit group results. Initial pRF size estimated by fMRI is outlined (1x), note that only a subset of all scale factors (9 out of 19) is displayed. All maps use the same color scale as shown on the right.

**Supplementary Figure S6.**
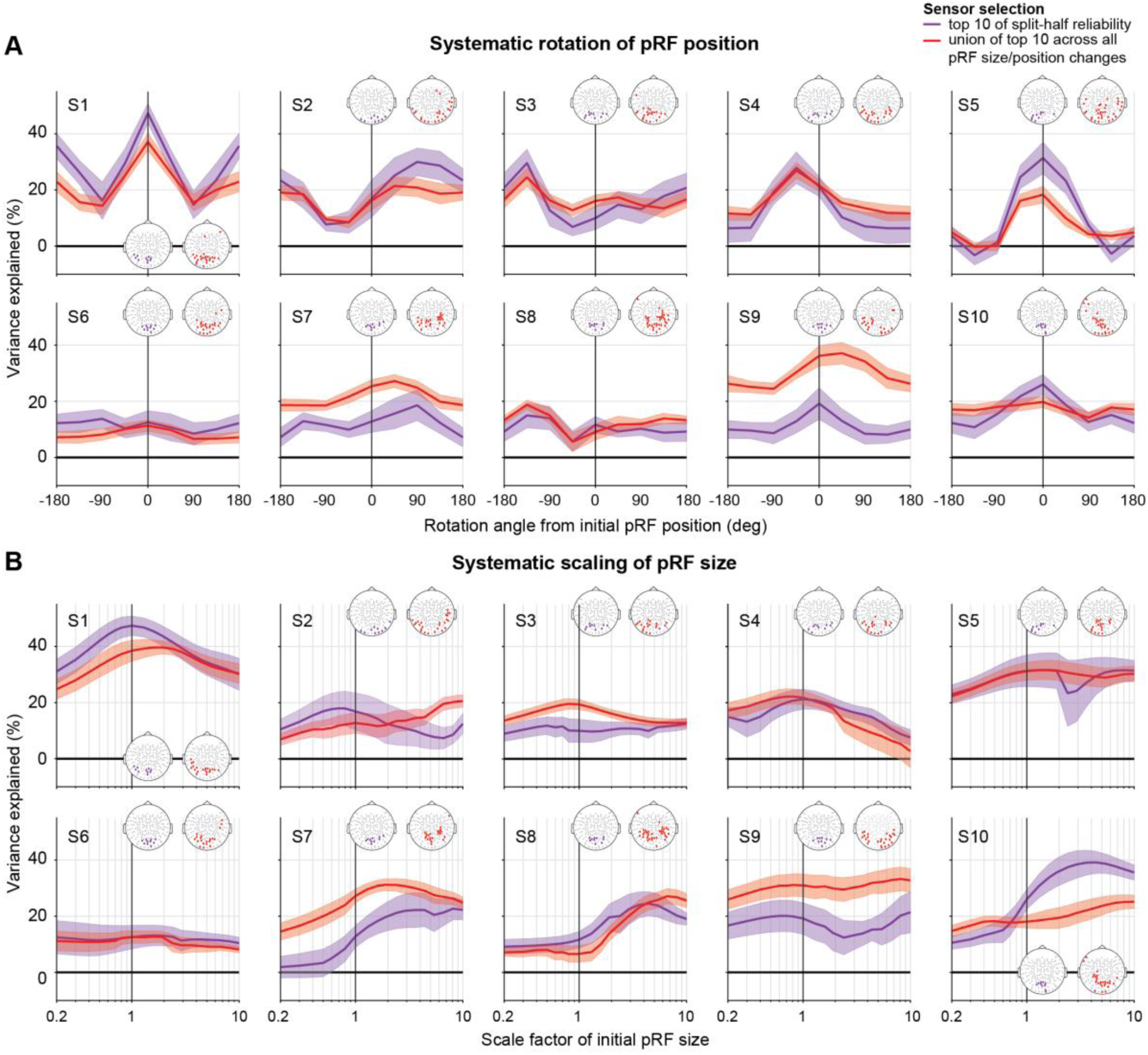
Effect of systematically varying pRF parameters using a data-based or model-based sensor selection in individual subjects. **(A) Variance explained by model when rotating pRF centers.** Red lines and shaded areas show sensor-wise average and ±1 standard error of the mean across the selected sensors using a model-based sensor selection (*i.e.*, compute the average in individual subjects from the union of top 10 sensors across all pRF position variations). Purple lines and shaded area show average and ±1 standard error of the mean across using a data-based sensor selection (*i.e.*, compute average in individual subjects from top 10 sensors of 10 Hz SSVEF split-half reliability map). **(B) Variance explained by model when scaling pRF size.** Same as (A) but now for pRF size scale factor variations. Both panels A and B use the same color scheme as A and B. Two schematic head plots show included sensors when using the union of top 10 sensors across all pRF position/size variations (red) or top 10 sensors of 10 Hz SSVEF split-half reliability map (purple).

**Supplementary Figure S7.**
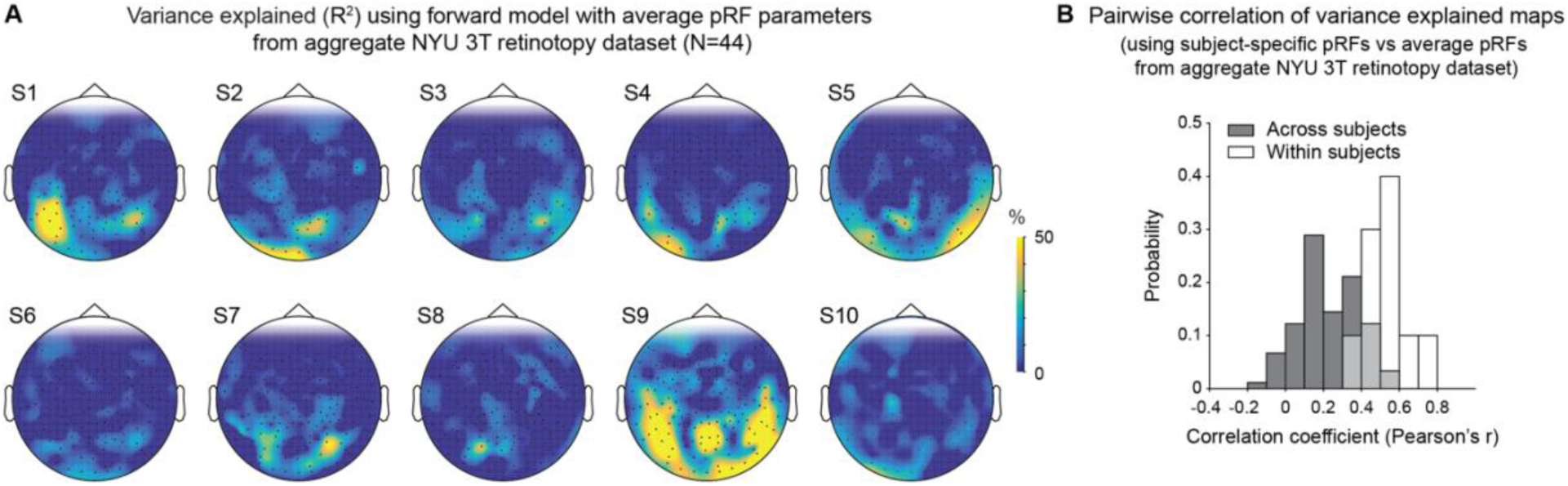
Variance explained by forward model using average pRF parameter maps from the aggregate NYU 3T retinotopy dataset (N=44). **(A) Topographic sensor maps of variance explained or all 10 individual subjects.** Variance explained by a forward model that uses average pRF maps from an aggregate retinotopy dataset collected at NYU’s 3T Prisma scanner (Himmelberg, Kurzawski, et al. (2021)). This aggregate data set uses a different stimulus but has a similar field-of-view and pRFs were analyzed with the same Vistasoft software (for more details on the aggregate retinotopy data, see Methods). Instead of using subject specific pRF maps to compute predicted MEG sensor responses, we used the average pRF maps projected onto individual subjects’ cortical surface. In some subjects (S1, S9) using the average pRF parameters from the NYU 3T retinotopy dataset explains up to a similar percent variance explained in individual sensor data as the subject specific pRF maps (see Supplementary Figure S1), whereas for other subjects the average pRF parameters result in lower variance explained by the model. **(B) Pairwise correlation of variance explained maps comparing forward model with subject-specific pRF maps to forward model with average pRF parameter maps from aggregate NYU 3T retinotopy dataset.** Probability of correlation coefficients is shown for comparisons within subjects (white bars), across subjects (dark gray bars), and overlap (light gray bars). Pairwise correlation coefficients within individual subjects are on average 0.6, which indicates that spatial topography maps of variance explained by the NYU 3T average pRF model are not identical to variance explained maps from the subject specific pRF model but overlap substantially. The within subject’s correlation is on average higher than the across subject’s correlation and shows that individual cortical curvature and head models play an important role in goodness of fit by our forward model.

## Notes

### Competing Interest Statement

The authors have declared no competing interest.

